# Reconstitution of 3’-processing of mammalian pre-mRNA reveals a central role of RBBP6

**DOI:** 10.1101/2021.11.02.466878

**Authors:** Moritz Schmidt, Florian Kluge, Felix Sandmeir, Uwe Kühn, Peter Schäfer, Christian Tüting, Christian Ihling, Elena Conti, Elmar Wahle

## Abstract

The 3’ ends of almost all eukaryotic mRNAs are generated in an essential two-step processing reaction, endonucleolytic cleavage of an extended precursor followed by the addition of a poly(A) tail. By reconstituting the reaction from overproduced and purified proteins, we provide a minimal list of fourteen polypeptides essential and two stimulatory for RNA processing. In a reaction depending on the polyadenylation signal AAUAAA, the reconstituted system cleaves pre-mRNA at a single preferred site corresponding to the one used *in vivo*. Among the proteins, Cleavage Factor I stimulates cleavage but is not essential, consistent with its prominent role in alternative polyadenylation. RBBP6 is required, structural data showing it to contact and presumably activate the endonuclease CPSF73 through its DWNN domain. The C-terminal domain of RNA polymerase II is dispensable. ATP, but not its hydrolysis, supports RNA cleavage by binding to the hClp1 subunit of cleavage factor II with submicromolar affinity.

## Introduction

The 3’ ends of eukaryotic mRNAs are generated by an essential co-transcriptional processing reaction: The nascent primary transcript is cleaved by an endonuclease, a poly(A) tail is added to the upstream cleavage fragment, and the downstream fragment is degraded. Pre-mRNA cleavage contributes to transcription termination by RNA polymerase II (pol II) (Eaton et al. 2020), and poly(A) tails are important in the initiation of translation (Jackson et al. 2010) and the control of mRNA half-life (Eisen et al. 2020). Alternative polyadenylation is common in higher eukaryotes and typically generates mRNAs with different regulatory sites in their 3’ UTRs (Gruber and Zavolan 2019). The only eukaryotic mRNAs not decorated with poly(A) tails are histone mRNAs in animal cells; their precursors are cleaved but not polyadenylated (Sun et al. 2020).

Cleavage and polyadenylation are carried out by a large protein complex (Xiang et al. 2014; Kumar et al. 2019) containing, in mammals, four heterooligomeric complexes: Cleavage and Polyadenylation Specificity Factor (CPSF), Cleavage Factors I and II (CF I and II), and Cleavage stimulation Factor (CstF). CPSF consists of two subcomplexes: The first, mammalian polyadenylation specificity factor (mPSF) (Schönemann et al. 2014; Clerici et al. 2018; Sun et al. 2018), contains CPSF160, WDR33, hFip1 and CPSF30 and recognizes the polyadenylation signal AAUAAA upstream of the cleavage site. Poly(A) polymerase (PAP) is not stably associated with mPSF, but relies on this factor in AAUAAA-dependent polyadenylation. PAP is also required for pre-mRNA cleavage. The second subcomplex of CPSF is mammalian cleavage factor (mCF) (Sun et al. 2020; Zhang et al. 2020), composed of the endonuclease CPSF73, the related but catalytically inactive protein CPSF100, symplekin, and perhaps CstF64 (Mandel et al. 2006; Sullivan et al. 2009; Sun et al. 2020; Yang et al. 2020; Zhang et al. 2020). CF I consists of a dimer of CF I-25 and two copies of CF I-59 or CF I-68 (Ruegsegger et al. 1998; Yang et al. 2011) and recognizes UGUA motifs upstream of AAUAAA (Venkataraman et al. 2005; Yang et al. 2011). CF II is a heterodimer of hPcf11 and hClp1 (Schäfer et al. 2018). Pcf11 binds the pol II C-terminal domain (CTD) and is important for the coupling of 3’ end processing to transcription termination (Meinhart and Cramer 2004; Kamieniarz-Gdula et al. 2019). Clp1 has an RNA 5’ kinase activity of unknown function (Weitzer et al. 2015). CstF, composed of two copies each of CstF77, CstF64, and CstF50 (Takagaki and Manley 1994; Yang et al. 2018), binds GU-rich downstream elements. In addition to these proteins, RBBP6 is likely involved, based on its sequence similarity to yeast 3’ processing factor Mpe1, its presence in affinity-purified 3’ processing complexes (Shi et al. 2009) and functional evidence (Di Giammartino et al. 2014). The proteins are represented schematically in **Fig. 1A**. In addition, the nuclear poly(A) binding protein (PABPN1) participates in poly(A) tail extension (Kühn et al. 2017), but may also play a role in poly(A) site choice (de Klerk et al. 2012; Jenal et al. 2012). Several other proteins have been suggested to be involved in 3’ end formation, the CTD of pol II among them (Hirose and Manley 1998).

**Fig. 1:**
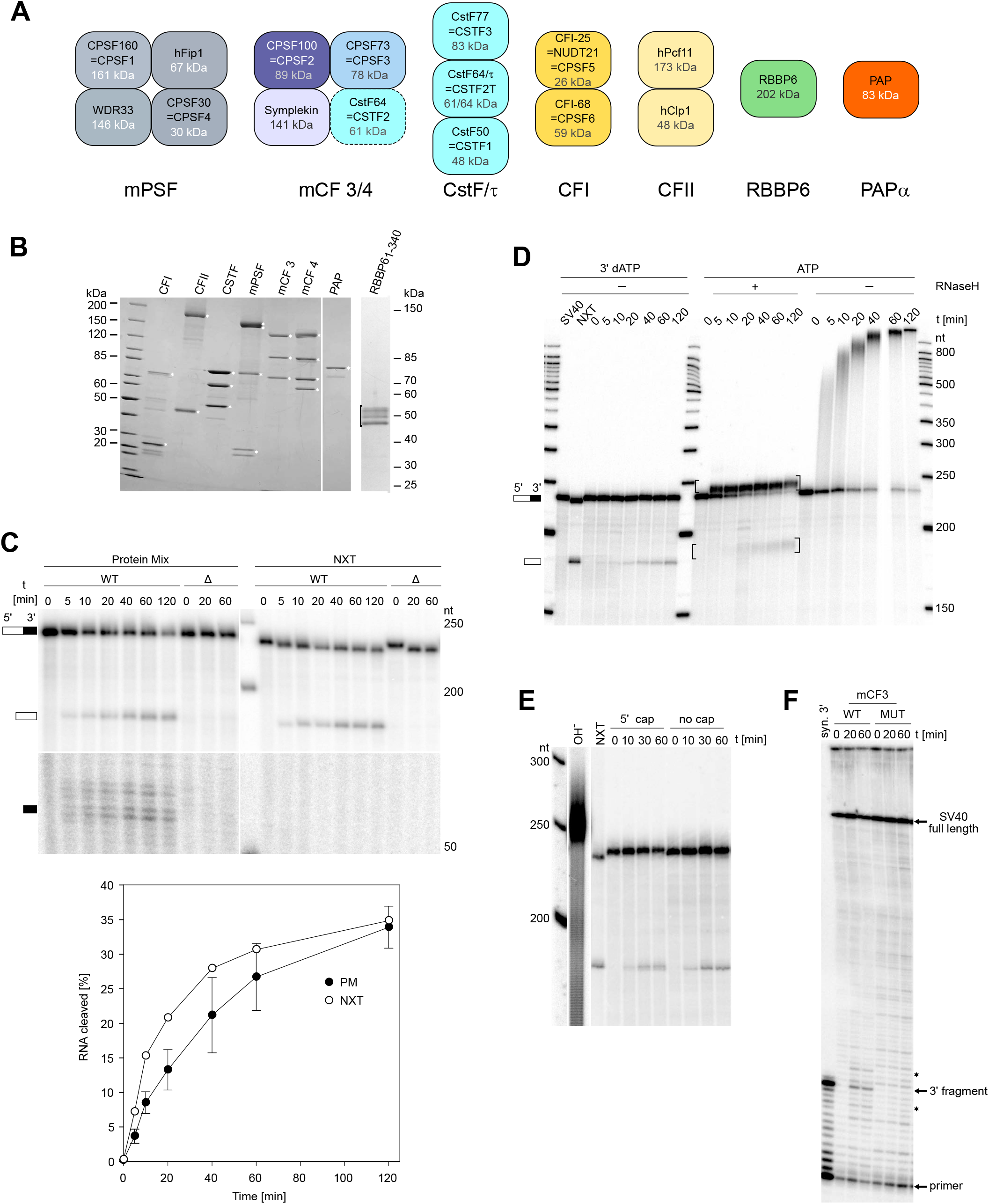
Reconstitution of pre-mRNA 3’ processing from sixteen purified polypeptides. (A) Schematic representation of proteins participating in pre-mRNA cleavage. (B) Proteins used in this study were separated by SDS-PAGE and stained with Coomassie. CPSF160 and WDR33 are not resolved, and CPSF30 runs as a doublet. Two versions of mCF are shown (see text). In the RBBP6 preparation, all major bands reacted with anti-RBBP6 in western blotting. (C) Kinetics of cleavage of the SV40 late RNA in the presence of WT PAP and 3’-dATP. Left, reaction with purified proteins. Right, reaction in nuclear extract. The substrate RNA (wild-type or mutant) is shortened by a few nucleotides from the 3’ end in nuclear extract. Downstream fragments are shown in a longer exposure of the bottom part of the same gel. Quantification in the bottom panel (n = 3). (D) Kinetics of cleavage and polyadenylation with purified proteins in the presence of 3’-dATP vs ATP. Only the upstream cleavage product is shown. Reactions were analyzed directly or after RNase H/oligo(dT) treatment. Brackets indicate digestion products with oligo(A) stubs. (E) Analysis of 5’ cleavage fragment. A reaction with uncapped RNA reveals a single 5’ product whereas capped RNA generates two products, presumably reflecting incomplete capping. Products co-migrate with the one generated in nuclear extract. A partial alkaline hydrolysis ladder of an end-labeled RNA (OH^−^) demonstrates single-nucleotide resolution. This lane was cut from the same gel, and contrast was enhanced to reveal individual bands. (F) Analysis of 3’ cleavage fragment. Reactions were carried out with unlabeled substrate RNA and either wild-type (WT) mCF3 or mCF3 containing inactive CPSF73 (MUT). The cleavage site was mapped by primer extension. Products corresponding to uncleaved RNA and the major 3’ cleavage product are indicated. Minor products are labeled with asterisks. For the first lane, a synthetic 3’ cleavage fragment was used as template.

3’ end processing of yeast mRNA precursors has been reconstituted from a set of proteins similar but not identical to the mammalian factors (Hill et al. 2019) (**Table S1**). 3’ end cleavage of mammalian histone mRNAs has also been reconstituted; required factors include mCF described above (Sun et al. 2020). In contrast, reconstitution of mammalian 3’ end processing has been limited to the second step, polyadenylation, which requires mPSF, PAP and PABPN1 (Schönemann et al. 2014). Investigations of the cleavage reaction have relied on combinations of ‘native’ proteins purified to various degrees or on the use of unfractionated nuclear extract. Thus, the list of proteins essential for the reaction is not known with certainty, and even the involvement of ATP and perhaps other small molecules has been controversial. To clarify these issues, we have reconstituted mammalian pre-mRNA cleavage from recombinant proteins.

## Results

### Reconstitution of pre-mRNA cleavage

For reconstitution of the pre-mRNA 3’ processing reaction, established and putative mammalian 3’ processing factors were overproduced and purified (**Fig. 1B; Suppl. Fig. 2D**). A mixture of mPSF, mCF, CF I, CF II, CstF, PAP, and RBBP6, in the presence of 3’-dATP, cleaved an RNA containing the SV40 late 3’ processing signal into an upstream fragment of the expected size plus a heterogeneous set of downstream fragments (**Fig. 1C)**. A point mutation in AAUAAA prevented cleavage. The reaction proceeded without a pronounced lag phase at a slowly decreasing rate. The efficiency was variable; 30 – 40% of the precursor were cleaved under optimal conditions. Cleavage proceeded similarly in nuclear extract, except that the downstream fragment was degraded and the precursor RNA was slightly shortened from its 3’ end (**Fig. 1C**). In these experiments, chain-terminating 3’-dATP prevented polyadenylation of the RNA to allow direct detection of the upstream cleavage product. When the reaction was carried out with ATP instead, polyadenylation took place. RNase H/oligo(dT) digestion revealed polyadenylation both of the upstream cleavage product and of the precursor RNA, as reported for experiments in nuclear extract (Zarkower et al. 1986) (**Fig. 1D**).

The expected cleavage site of the SV40 late RNA is between two A residues in the second of three consecutive CAA repeats (Sheets et al. 1987) (www.ncbi.nlm.nih.gov/genbank; entry J02400.1). The cleavage site in the reconstituted reaction was first analyzed by high resolution gel electrophoresis of the upstream cleavage fragment: Use of an uncapped RNA to avoid heterogeneity due to incomplete cap incorporation demonstrated dominant cleavage at a single site (**Fig. 1E**). Thus, heterogeneity of the downstream fragment could be the result of transcript heterogeneity at the 3’ end and/or of limited 5’ exonuclease digestion subsequent to cleavage. To avoid complications by potential 3’ end heterogeneity, we analyzed 5’ ends of the downstream fragments by primer extension. A chemically synthesized RNA representing the expected downstream fragment (Sheets et al. 1987) served as a positive control. The dominant reverse transcription product obtained from authentic downstream fragments corresponded to the product obtained from the synthetic RNA (**Fig. 1F**). Most other bands in the neighborhood were background, not being affected by an active site mutation in CPSF73. However, two bands, three nucleotides upstream and downstream, respectively, of the main product, were also dependent on CPSF73 activity. Presumably, they represent minor cleavage sites in either of the two neighboring CAA repeats. Together, the data show that cleavage occurs predominantly at the site mapped earlier. In contrast, the reconstituted yeast system cleaved a model substrate RNA with equal probability at any of three adjacent phosphodiester bonds (Hill et al. 2019).

In several independent experiments, average recovery of the downstream fragment was only ~10% of the upstream fragment. Thus, most of the downstream fragment is degraded. However, when the synthetic downstream fragment, carrying a 5’-monophosphate like the authentic fragment, was 3’-labeled and incubated with the mixture of all cleavage factors, no significant degradation took place (**Suppl. Fig. 1**), suggesting that degradation of the 3’ cleavage fragment only occurs in the context of the processing reaction.

### Definition of a minimal cleavage complex: CF I is not essential

We used the reconstitution system to define the minimal set of factors essential for pre-mRNA cleavage. Most factors were indispensable; their individual omission abolished the reaction (**Fig. 2A**). PAP was among the essential proteins, but its catalytic activity was not required for RNA cleavage (**Suppl. Fig. 2A**). The only protein that could be omitted without complete loss of cleavage activity was CF I (**Fig. 2A**). A titration experiment confirmed that CF I was stimulatory for cleavage of the SV40 late RNA but not essential (**Fig. 2B**). Whereas the SV40 late RNA has a single copy of the CF I binding motif, UGUA, adenovirus L3 contains the preferred configuration (Yang et al. 2011), two copies upstream of AAUAAA. CF I stimulated cleavage of wild-type L3 RNA. As expected, the effect was strongly reduced by mutation of the upstream copies of UGUA to UcUA, with or without mutation of a third copy immediately upstream of the cleavage site (**Fig. 2B)**. Importantly, the mutations had no significant effect on the efficiency of cleavage when no CF I was added. Thus, there was no functionally relevant amount of CF I present as a contamination. CF I has been reported also to stimulate polyadenylation (Brown and Gilmartin 2003), but in our hands the factor had no detectable effect on the polyadenylation of a ‘pre-cleaved’ L3 RNA even when mPSF and PAP were present at limiting concentrations (**Suppl. Fig. 3**).

**Fig. 2:**
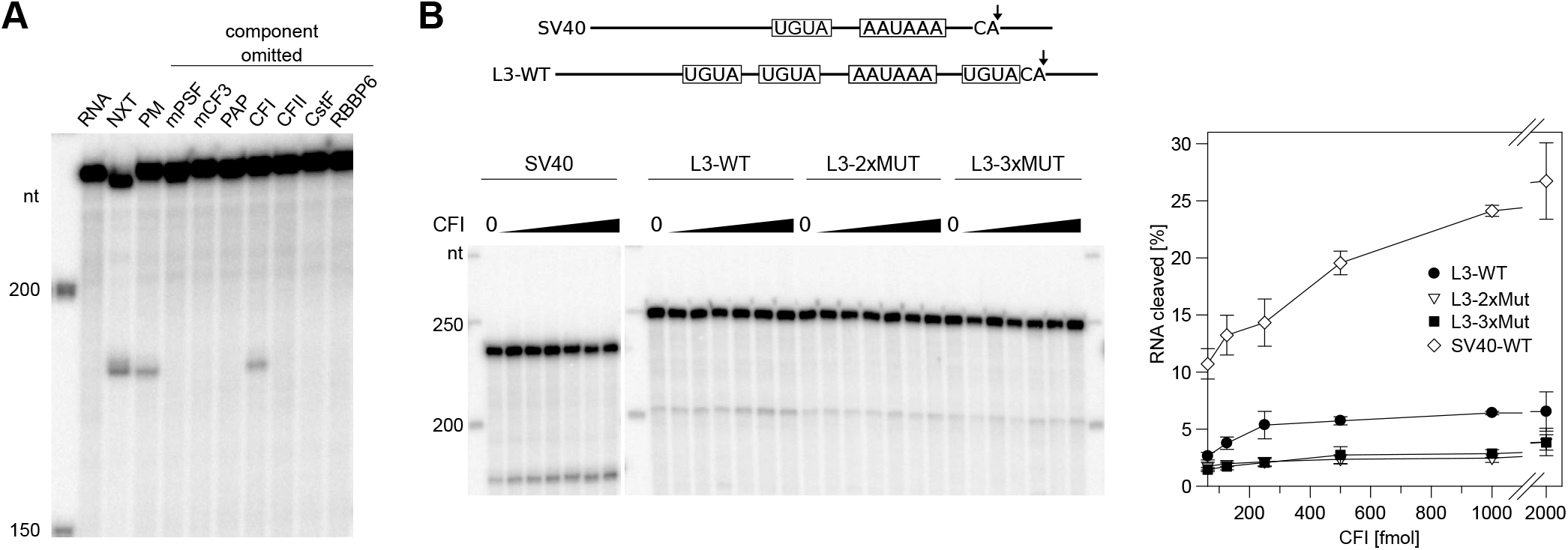
CF I is not an essential 3’ processing factor. (A) CF I is the only non-essential component of the reconstituted cleavage reaction. The SV40 late precursor RNA was processed in nuclear extract (NXT) or with a mixture of purified proteins (PM). Individual components listed at the top were left out. (B) Stimulation of cleavage by CF I depends on UGUA motifs. SV40 late and L3 RNAs with their UGUA motifs are sketched at the top. In L3-2xMUT and L3-3xMUT, the first two UGUA motifs or all three, respectively, were mutated. Titrations of CF I were carried out. Right, cleavage efficiencies (average of n = 3 +/− SD). Efficiencies without CF I addition were 2.2 ± 0.3% for L3, 1.8 ± 0.4% for L3-2xMUT and 1.9 ± 0.4% for L3-3xMUT.

Cleavage in the reconstituted system proceeded without addition of the pol II CTD, and MS analysis did not detect the large subunit of pol II as a contamination. The CTD was overproduced in *E. coli* or Sf21 cells and purified. The Sf21-made CTD was found to be phosphorylated, and phosphorylation of Ser2, important for CTD function in 3’ end processing *in vivo* (Ahn et al. 2004; Davidson et al. 2014), was detectable (**Suppl. Fig. 2B**). Still, the addition of either preparation to reconstituted cleavage reactions had only a small (~ 1.3fold) stimulatory effect of uncertain significance (**Suppl. Fig. 2C**).

Five additional proteins (PABPN1, SSU72, a protein phosphatase 1 complex, CDC73 and XRN2) potentially participating in the 3’ cleavage reaction had little effect on the reaction (**Suppl. Fig. 2D,E**).

### RBBP6 plays a central role in pre-mRNA cleavage

RBBP6 is essential for pre-mRNA cleavage **(Fig. 2A)**. At its N-terminus, the protein contains a ubiquitin-like or DWNN domain, a CCHC zinc knuckle and a RING domain (Pugh et al. 2006; Kappo et al. 2012), the rest consisting of low complexity/disordered regions (**Fig. 3A**). In agreement with (Di Giammartino et al. 2014), N-terminal fragments were active in pre-mRNA cleavage: Fragments containing the first two structured domains (amino acids 1-253) had a similar activity as longer fragments (aa 1-340 or 1-780), but full-length protein was active at lower concentrations (**Fig. 3A,B**). Inactivity of the DWNN domain alone (aa 1-81) (**data not shown**) suggested a requirement for the zinc knuckle. Accordingly, a C161G/C164G double mutation in this domain inactivated RBBP6 (**Fig. 3C**), consistent with observations in yeast (Lee and Moore 2014). The shortest active fragment was lacking the RING domain, and an internal deletion of the domain (aa 253-326) from full-length RBBP6 was tolerated (**Fig. 3A,B**). Thus, a potential E3 ligase function is not essential for the role of RBBP6 in RNA cleavage. The RING domain dimerizes at high concentrations (Kappo et al. 2012). However, the RING domain is dispensable, and a point mutation inhibiting dimerization (N312D) (Kappo et al. 2012) did not inactivate RBBP6_1-780_ (**Fig. 3C**). Thus, dimerization is not important for cleavage. As also reported for Mpe1 (Lee and Moore 2014), RBBP6_1-780_ bound RNA (**Fig. 3D,E**).

**Fig. 3:**
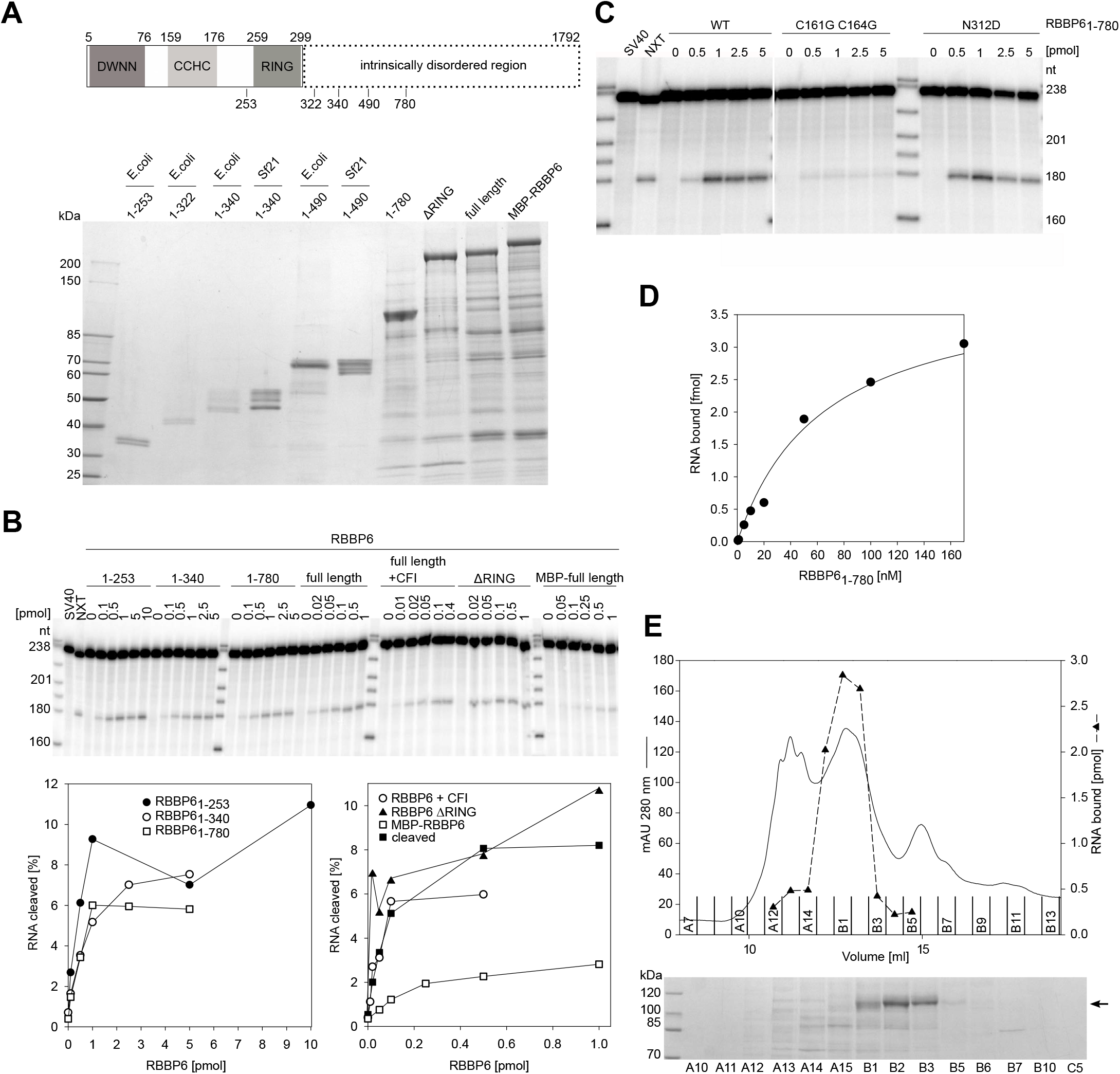
Two structured domains of RBBP6 are sufficient for pre-mRNA cleavage. (A) Top panel: Domain structure of RBBP6. Domain boundaries are indicated at the top and deletion boundaries at the bottom. Bottom panel: SDS polyacrylamide gel analysis of RBBP6 preparations. (B) Titration of RBBP6 variants in cleavage assays. ‘Full-length’ was MBP-RBBP6 with the MBP tag cleaved off; ‘full-length plus CF I’ was RBBP6 from a co-expression with CF I. A quantification is shown at the bottom. (C) Cleavage activity of RBBP6_1-780_ carrying point mutations. (D) RBBP6_1-780_ was titrated in a filter-binding assay with SV40 late RNA (compare Supplemental Table S2). (E) RNA binding activity co-purifies with RBBP6. Top panel, UV profile and RNA binding activity of the final Resource Q column of an RBBP6_1-780_ purification. Bottom panel, SDS-PAGE of the same fractions. The main RBBP6 band is marked.

The experiments suggested that RBBP6 might have a role in mammalian pre-mRNA cleavage analogous to the one of Mpe1 in yeast, helping to activate the CPSF73 endonuclease in the context of a correct pre-mRNA cleavage site. Yeast Mpe1 interacts directly with Ysh1, the orthologue of CPSF73 (Hill et al. 2019), and shares significant sequence similarity with the N-terminal part of RBBP6. We therefore tested whether RBBP6_1-335_, which includes the three structured domains and provides RBBP6 function in RNA cleavage, may engage CPSF73 in an interaction similar to Mpe1-Ysh1. In pull-down experiments using purified proteins, RBBP6_1-335_ efficiently co-purified with a seven-subunit CPSF complex, even without addition of an RNA substrate (**Fig. 4A, lane 5**). Interestingly, we observed a reproducible, albeit weaker interaction with a four-subunit mCF alone (**Fig. 4A, lane 4**). Based on these results, we reconstituted an eight-subunit CPSF-RBBP6_1-335_ complex for single-particle cryo-EM analysis.

**Fig. 4:**
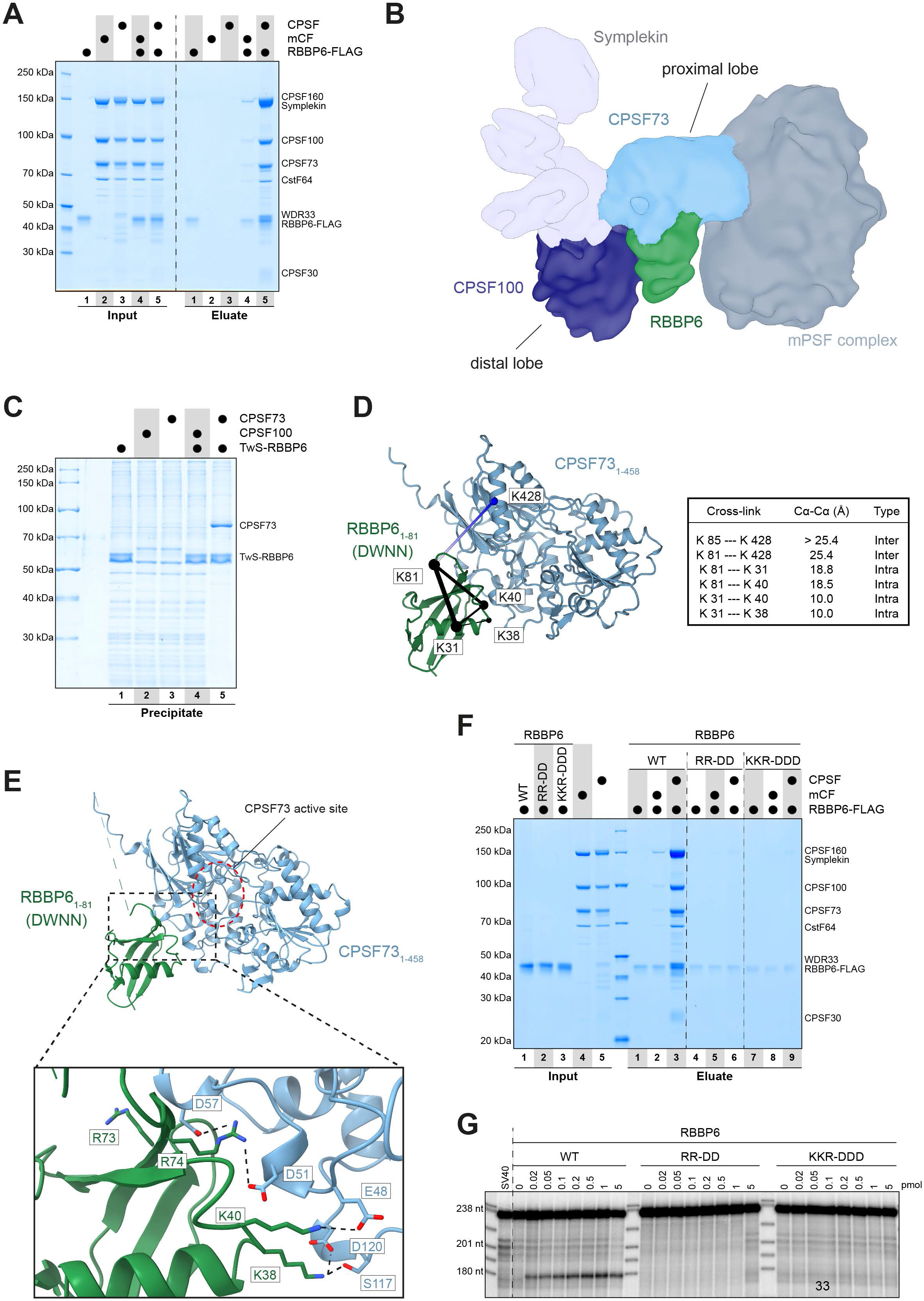
RBBP6 contacts the ß-lactamase domain of CPSF73. (A) Coomassie-stained SDS-PAGE analysis of pull-down experiment showing that RBBP6 directly interacts with CPSF and, albeit weaker, mCF. (B) Filtered and segmented cryo-EM map of the eight-subunit CPSF-RBBP6 complex. The proximal and distal lobes of mCF are indicated. (C) Coomassie-stained SDS-PAGE analysis of pull-down experiment showing that RBBP6 directly interacts with CPSF73 and not CPSF100. (D) Cross-links between CPSF73_1-458_ and RBBP6_1-81_ mapped onto the computationally predicted model of the complex. Intra- and intermolecular cross-links are in black and blue, respectively, with the thickness of the line indicating their score (thicker = higher score), and the respective residues indicated as spheres. A cross-link to an unmodeled region is shown in white (placed to the closest visible Cα atom). The table lists measured distances for visualized cross-links. (E) Computationally predicted structural model of the CPSF73_1-458_-RBBP6_1-81_ complex with close-up of the putative binding interface. The CPSF73 active site is indicated. Possible ionic interactions are shown with black dotted lines. Labeled positively charged RBBP6 residues were changed to aspartic acid. (F) Coomassie-stained SDS-PAGE analysis of pull-down experiment showing that reverse-charged mutations in the predicted RBBP6-CPSF73 interface disrupt the interaction of RBBP6 with mCF and CPSF. (G) RBBP6_1-335_, either wild-type or containing the same mutations as in (F), was titrated into cleavage reactions.

The 2D class averages clearly showed structural details, but preferred orientation and inherent flexibility limited the resolution of the 3D reconstruction (**Fig. 4B**; **Suppl. Fig. 4**). The resulting cryo-EM map, filtered to 10 Å, showed similar features as the previously reported reconstruction of CPSF in isolation (**Suppl. Fig. 4D**) (Zhang et al. 2020). The assembly contained a rigid mPSF core adjacent to a cloverleaf structure with two globular lobes (corresponding to CPSF100 and CPSF73) and an extended lobe (corresponding to symplekin). The proximal lobe, which directly interacts with mPSF, could not be assigned with confidence to CPSF73 or CPSF100 in earlier work (Zhang et al. 2020). In our cryo-EM reconstruction, extra density was protruding from the proximal lobe, suggesting this density may correspond to the additional subunit present in the sample, RBBP6_1-335_ (**Suppl. Fig. 4D,E**), and in turn pointing to the proximal lobe as corresponding to CPSF73. To test this hypothesis, we overexpressed RBBP6_1-335_ with either CPSF100 or CPSF73 in HEK cells and performed co-precipitation experiments. RBBP6_1-335_ indeed reproducibly co-purified with CPSF73 and not with CPSF100 (**Fig. 4C**).

Cross-linking mass spectrometry (XL-MS) was used to examine the structural arrangement in the context of the RBBP6-bound CPSF complex. After treatment with Bis(sulfosuccinimidyl)suberate (BS3), cross-links between the RBBP6 DWNN domain and CPSF73 were repeatedly detected (**Suppl. Fig. 5A**). Based on these data, a structural model of the CPSF73-RBBP6 complex was predicted with the help of AlphaFold2 (**Suppl. Fig. 5B**) (Jumper et al. 2021; Mirdita et al. 2021). In the model, the RBBP6 DWNN domain docks onto the metallo-ß-lactamase domain of CPSF73 in close proximity to the active site opening (**Fig. 4E**), similar to the binding observed for the yeast orthologues (**Suppl. Fig. 5C**) (Hill et al. 2019). Mapping the cross-links onto the model revealed that they were well within the expected distance range of BS3 (**Fig. 4D**). The CPSF73-RBBP6 model fitted into the size and shape of the proximal lobe and features of the protruding density. The protruding density extends further, and additional cross-links were identified between the region of RBBP6 downstream of the DWNN domain and CPSF100, WDR33 and symplekin (**Suppl. Fig. 5A**). This suggested additional contacts in this more flexible part, consistent with the relative strengths of the interactions observed in the pull-down assays (**Fig. 4A**).

To test our model, we introduced mutations aimed at disrupting the interaction between RBBP6 and CPSF73: Positively charged residues in the CPSF73-binding interface of RBBP6_1-335_ were changed to aspartic acid (**Fig. 4E**). These mutations disrupted the interaction in pull-down experiments using purified proteins as well as in co-precipitation experiments in HEK cell lysates (**Fig. 4F**; **Suppl. Fig. 5D**) and obliterated the activity of RBBP6_1-335_ in pre-mRNA cleavage (**Fig. 4G**). Thus, the identified interface between the RBBP6 DWNN domain and CPSF73 is essential for recruiting RBBP6 to the CPSF complex prior to RNA cleavage. At the same time, the data reveal that the CPSF73 subunit is adjacent to the mPSF core and that the RBBP6-CPSF73 interaction is indeed conserved from yeast to human.

### Subunits of CstF and mCF active in pre-mRNA cleavage

Reconstituted cleavage reactions were further used to define the compositions of active CstF and mCF. We prepared CstF containing only CstF64 [CstF (64/64)] or its paralog 64τ [CstF (τ/τ)] or a mixture of both [CstF (64/τ)] (**Fig. 5A**). The existence of mixed complexes was demonstrated by the ability of FLAG-tagged CstF64τ to pull down untagged CstF64 together with CstF77 and CstF50 (**Fig. 5A**, right panel). All three versions of the complex supported cleavage (**Fig. 5B**), consistent with studies suggesting overlapping functions of CstF64 and 64τ (Ruepp et al. 2011; Yao et al. 2013). A slightly lower activity of τ-containing complexes (**Fig. 5B**; **Suppl. Fig. 6A; Table S2**) was repeatedly observed with independent preparations. In addition to CF I, the CstF subunit CstF50 is the only component in mammalian 3’ end processing that has no yeast ortholog. Partial CstF complexes lacking CstF50 (CstFΔ50 or CstFτΔ50) were nearly inactive. Complementation with individually purified CstF50 restored the activity (**Fig. 5A, B**), showing that the subunit is important for pre-mRNA cleavage. CstF64 or CstF50 alone or in combination were inactive, suggesting an essential role of CstF77 (**Fig. 5A, B**). In this case, complementation was not possible since we were unable to purify CstF77 by itself.

**Fig. 5:**
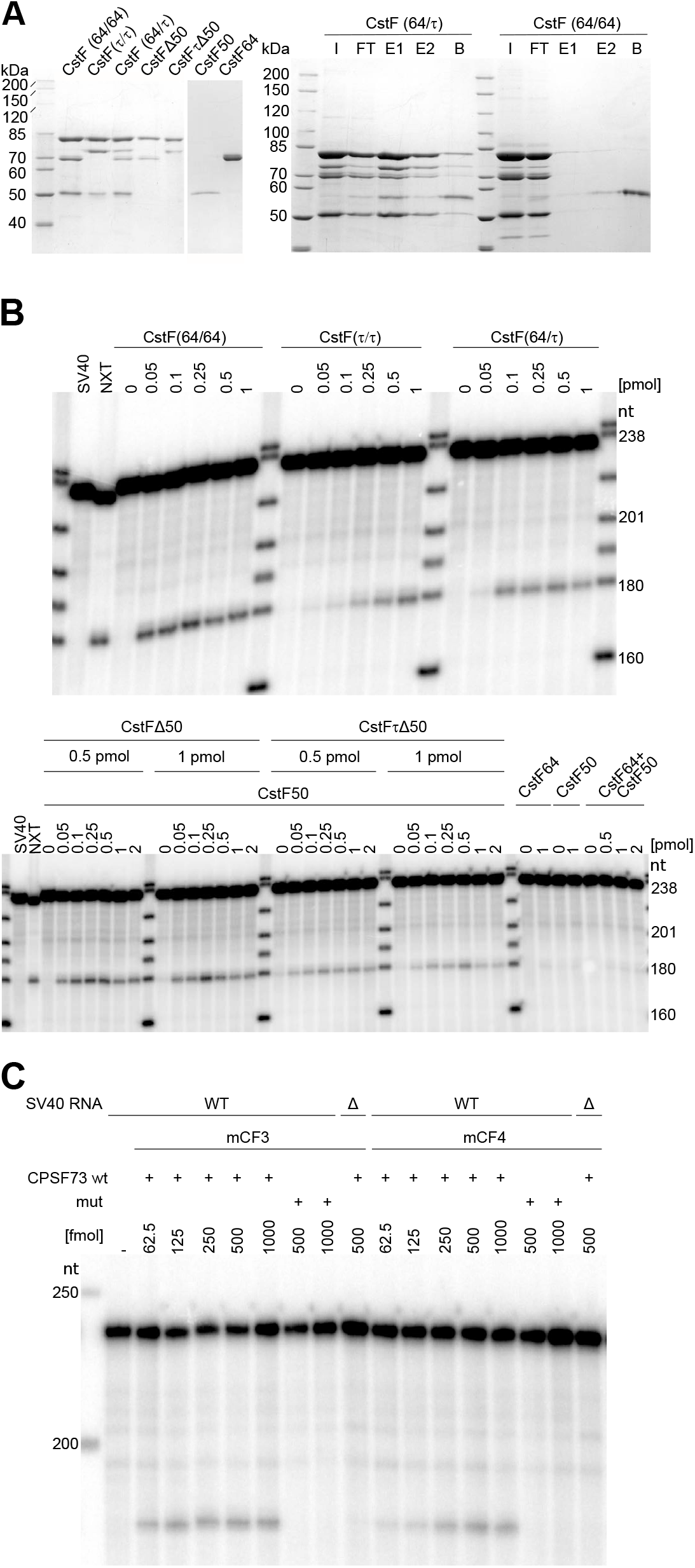
Composition of CstF and mCF active in pre-mRNA cleavage. (A) Left, different versions of CstF and its subunits were analyzed by SDS PAGE and Coomassie staining. Right, CstF (64/τ) with a FLAG tag on CstF64τ and, as a control, CstF (64/64) without a FLAG tag were used in a FLAG pull-down experiment. I, input; FT, flow-through; E1, E2, FLAG peptide eluates; B, material remaining on the beads eluted with SDS sample buffer. Proteins were detected by Coomassie staining. The identity of the band running at the CstF64τ position in CstF (64/64) is unknown; based on western blots, it is neither 64τ nor a fragment of CstF77. (B) Both CstF64 and 64τ are active in pre-mRNA cleavage, and CstF50 is essential. Proteins shown in (A) and their combinations were tested in cleavage assays. (C) Both mCF3 and mCF4 function in pre-mRNA cleavage. The two protein complexes, each with wild-type (+) or point-mutated (−) CPSF73, were tested in cleavage assays. The SV40 late Δ RNA served as negative control.

mCF could be prepared from three or four subunits (mCF3 or mCF4, respectively), containing CstF64 or not (**Fig. 1A**). Both preparations supported pre-mRNA cleavage (**Fig. 5C**). Thus, mCF lacking CstF64 is fully competent for pre-mRNA cleavage, as also observed in histone 3’ processing (Sun et al. 2020; Yang et al. 2020). In XL-MS experiments employing the amine-reactive cross-linker DSBU (Muller et al. 2010) in the context of a mCF3-containing cleavage reaction, the dominant partners of CstF64 were the other two CstF subunits, but cross-links to the mCF subunit symplekin were also reproducibly observed (**Suppl. Fig. 7)**, suggesting that the CstF64 – symplekin interaction (Takagaki and Manley 2000; Ruepp et al. 2011) can occur under reaction conditions. As expected, a double point mutation in the active site of CPSF73 (D75K, H76A) (Mandel et al. 2006) abolished cleavage activity (**Figs. 1F, 5C**). mCF3 bound the SV40 late RNA with a K_50_ of ~3 nM, barely different from mCFI (**Suppl. Fig. 6B,C; Table S2**).

DSBU-mediated cross-linking also suggested the existence of novel homodimers in the cleavage complex: Homotypic cross-links of hFip1 were consistent with the presence of more than one copy in CPSF (Hamilton and Tong 2020) and CPF (Casanal et al. 2017). Several homotypic cross-links seen for both subunits of CF II (**Suppl. Fig. 7**) suggested the formation of at least a tetramer under reaction conditions, whereas the isolated factor behaves like a heterodimer (Schäfer et al. 2018).

### Pre-mRNA cleavage is ATP-dependent

Experiments in nuclear extract initially suggested that ATP was required for RNA cleavage (Moore and Sharp 1985). In contrast, a later study concluded that ATP was dispensable, but that creatine phosphate, originally added to replenish ATP degraded by ATPases in the nuclear extract, played an essential role independent of ATP regeneration (Hirose and Manley 1997).

Neither creatine phosphate nor creatine kinase were included in our reconstitution assays, and their addition had no effect as compared to several control compounds (**Fig. 6A**).

**Fig. 6:**
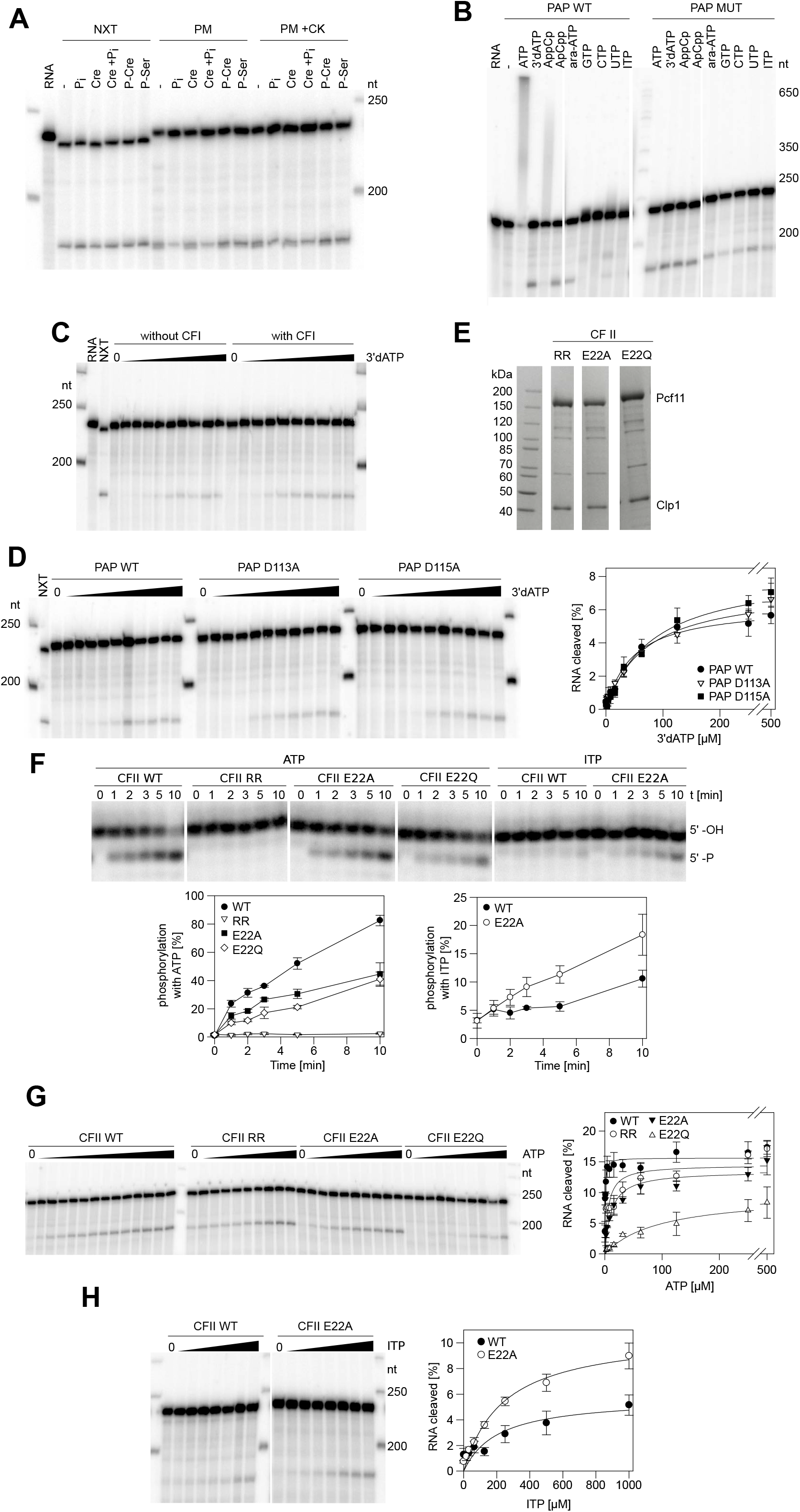
Pre-mRNA cleavage depends on ATP. (A) Creatine phosphate is not required for cleavage. Cleavage reactions were carried out with nuclear extract (NXT), a mixture of purified proteins (PM) or the same mixture plus creatine kinase (5 μg/ml). In the presence of 0.5 mM 3’-dATP, reactions were supplemented with inorganic phosphate (pH 8.0), creatine, a mixture of both, phosphocreatine or phosphoserine as indicated, each at 20 mM. (B) ATP is essential for cleavage, but cleavage of phosphoanhydride bonds is not. Reactions contained the nucleotides indicated, each at 0.5 mM, and either WT PAP or PAP D115A. With WT PAP, cleavage in the presence of CTP, GTP or ITP was not visible, presumably due to heterogeneous limited extension of the cleavage product. (C) A cleavage reaction lacking CF I is still ATP-dependent. Reactions were carried out with or without CF I as indicated. 3’-dATP was titrated between 3.9 and 1000 μM. (D) Mutations in the active site of PAP do not affect the ATP dependence of RNA cleavage. Reactions contained WT PAP or mutants as indicated. 3’-dATP was titrated between 2 and 1000 μM. Right, average of n = 3, highest ATP concentration omitted. (E) Mutant hClp1 forms a stable complex with hPCF11. The Coomassie-stained SDS PAGE shows the peak fraction from each Mono Q column. Based on western analysis of comparable wild-type preparations, most additional bands are breakdown products of hPcf11. (F) utations in the ATP binding site of hClp1 affect the RNA 5’ kinase activity of CF II. CF II preparations (**E**) were tested in kinase assays with ATP or ITP. Top, representative time courses. Bottom, average of n = 3. (G) Mutations in the ATP binding site of hClp1 affect ATP dependence of RNA cleavage. CF II preparations (**E**) were tested in cleavage assays with mutant PAP. ATP was titrated from 0.12 μM (WT) or 1.95 μM (mutants) to 500 μM. Top, representative experiment. Bottom, average of n = 3. (H) A mutation in the ATP binding site of hClp1 changes the nucleotide specificity of cleavage. The proteins indicated were tested as in (**G**) except that ITP was titrated. Top, representative experiment. Bottom, average of n = 3.

The ATP-dependence of cleavage in the reconstituted reaction was assayed in the presence of either wild-type PAP or catalytically disabled PAP D115A. Cleavage was clearly dependent on the addition of ATP or one of several analogs (**Fig. 6B,** compare minus ATP lane to other lanes). In the presence of ATP, both the 5’ cleavage product and the substrate RNA were polyadenylated with WT PAP as above, but use of the mutant revealed cleavage. Methyleno-ATP analogs with non-cleavable α-β or β-γ bonds (APCPP and APPCP, respectively) supported RNA cleavage, in agreement with results obtained in nuclear extract (Moore and Sharp 1985). APPCP also served as a substrate for poly(A) polymerase (Bienroth et al. 1993). Activity of the methyleno analogs eliminates the possibility that the essential role of ATP is to serve as a substrate for PAP or the RNA kinase activity of hClp1 and agrees with the fact that mutational inactivation of either enzyme does not prevent cleavage (**Suppl. Fig. 2A**) (Schäfer et al. 2018). Arabino-ATP behaved like 3’-dATP (**Fig. 6B)**. At the 0.5 mM concentration used, ~1000fold above the K_A_ for ATP (see below), GTP, CTP, UTP and ITP supported cleavage to a significant extent.

Three constituents of the cleavage complex are known to bind ATP: CF I-25 (Coseno et al. 2008), PAP and hClp1. CF I-25 is not responsible for the ATP-dependence of cleavage, since ATP is essential whereas CF I is not. Importantly, a cleavage reaction lacking CF I was still ATP-dependent, ruling out CF I-25 as the relevant ATP binder (**Fig. 6C)**. While use of ATP as a substrate by PAP or hClp1 cannot explain the ATP dependence of cleavage, the nucleotide might act as an allosteric effector for either enzyme. Therefore, mutations in their ATP binding sites were tested. Amino acids D113 and D115 in PAP contact Mg^2+^-ATP, and mutants D113A and D115A reduce the enzyme’s affinity for ATP (Martin et al. 1999; Martin et al. 2000). If ATP binding to PAP were responsible for the ATP-dependence of cleavage, the mutations should result in a requirement for higher ATP concentrations in the cleavage reaction. However, the K_A_ values (concentrations of 3’-dATP that resulted in half-maximal activation of cleavage) were 46.3 ± 7.4 μM with wild-type PAP, 69.3 ± 10.6 μM with the D113A mutant and 81.5 ± 10.2 μM with D115A (**Fig. 6D**). This minor change in concentration dependence makes it unlikely that ATP binding to the PAP active site is responsible for the ATP-dependence of cleavage.

Based on a comparison to *C. elegans* Clp1 (Dikfidan et al. 2014), amino acids R288 and R293 of human Clp1 bind the γ-phosphate of ATP, whereas E22 forms a hydrogen bond to the 6-amino group of the base. Clp1 variants with mutations in these positions were purified as complexes with hPcf11 (**Fig. 6E**) and first tested for their RNA kinase activity. The R288A, R293K double mutant (Dikfidan et al. 2014) was inactive, and activities of the E22A and E22Q mutants were approximately twofold reduced under standard reaction conditions. ITP is lacking the 6-amino group hydrogen bonding to E22, and the E22A mutation facilitated the use of ITP in the kinase reaction (**Fig. 6F**). The same mutations were then tested, in the presence of PAP D115A, for the ATP dependence of RNA cleavage: The K_A_ value was 0.5 ± 0.1 μM ATP with wild-type hClp1, ~hundredfold lower than the K_A_ of 3’-dATP. The RR mutant reproducibly showed significant cleavage activity without ATP addition; the K_A_ was 5.7 ± 3.4 μM ATP. The E22Q mutant had reduced cleavage activity at all ATP concentrations tested; the K_A_ was 95.9 ± 19.8 μM. K_A_ with the E22A mutant was 10.9 ± 2.7 μM ATP (**Fig. 6G**). The E22A mutation also improved cleavage in the presence of ITP, in parallel with its effect in the kinase reaction **(Fig. 6H).**The fact that mutations in the ATP binding site of hClp1 alter the ATP concentration dependence and, less dramatically, the nucleotide specificity of pre-mRNA cleavage strongly suggests that ATP facilitates cleavage by serving as an allosteric effector of hClp1.

## Discussion

By reconstituting mammalian pre-mRNA 3’ processing from recombinant proteins, we have provided a minimal parts list for the reaction. The following components are necessary and sufficient for the reaction: mPSF, mCF, CstF, CF II, RBBP6 and PAP, fourteen polypeptides in total. PABPN1 is involved in polyadenylation, but not essential for pre-mRNA cleavage. The list of required factors closely matches the equivalent system from *S. cerevisiae* (Hill et al. 2019): Mammalian CstF50 and yeast Hrp1 (CF IB) are the only essential 3’ processing factors unique to their respective systems. In addition, yeast Pta1 is part of the phosphatase module of CPF, which is dispensable for 3’ processing (Nedea et al. 2003; Casanal et al. 2017; Hill et al. 2019), whereas the mammalian orthologue, symplekin, is part of mCF and presumably essential for cleavage. Orthologues to other subunits of the yeast phosphatase module are not required. The reconstituted mammalian system catalyzes cleavage of predominantly one phosphodiester bond at the expected position. The selection of the exact cleavage site seems to be more precise in comparison to the yeast system (Hill et al. 2019).

Based on experiments employing native factors purified from nuclear extract, CF I was considered an essential 3’ processing factor (Takagaki et al. 1989; Ruegsegger et al. 1998). Our experiments now show that CF I, which has no equivalent in *S. cerevisiae*, stimulates 3’ processing, but is not essential. CF I also binds more weakly to RNA than any of the other 3’ processing factors (**Table S2**). Both observations suggest that CF I selectively participates in the processing of some RNAs but not others. This is consistent with the prominent role of the protein in alternative polyadenylation (Gruber and Zavolan 2019): Reduced expression of CF I had a much stronger effect on alternative polyadenylation than the depletion of any other individual 3’ processing factor and consistently caused pronounced shifts to the use of proximal (upstream) polyadenylation sites in many genes. The CF I binding motif, UGUA, was enriched in distal sites neglected and depleted in proximal sites favored upon knock-down, consistent with preferred cross-linking of CF I to distal sites (Martin et al. 2012; Masamha et al. 2014; Li et al. 2015; Zhu et al. 2018). These data are explained by the model proposed by (Zhu et al. 2018): CF I is an activator of 3’ processing dependent on its binding site, UGUA. Reduced availability of CF I leads to a decreased use of UGUA-containing (distal) poly(A) sites, whereas proximal sites lacking UGUA make no use of CF I anyway and, thus, are not affected by its loss; consequently, their relative use increases. The model assumes that CF I is not essential for 3’ processing; this tacit assumption is now confirmed by our data.

RBBP6 was not identified in the original resolution and reconstitution experiments that uncovered the roles of all other 3’ processing factors. A proposed role in 3’ processing (Shi et al. 2009; Di Giammartino et al. 2014) is confirmed by our experiments. The DWNN and zinc knuckle domains suffice for RBBP6 function; the RING domain and, thus, a potential ubiquitin ligase activity is not essential. Activation of the endonuclease activity of CPSF73 must be tightly coupled to its incorporation into a 3’ processing complex assembled on the correct RNA site. Activation very likely involves a substantial structural rearrangement to open the active site for accommodation of a substrate RNA, as seen for the histone cleavage complex (Sun et al. 2020). As the RBBP6 DWNN domain is bound near the active site of CPSF73, we speculate that it might help to induce this structural change or might stabilize an open conformation (**Fig. 7**). Similarly, yeast Mpe1 has recently been proposed to couple endonuclease activation to poly(A) site recognition (Rodriguez-Molina et al. 2021).

**Fig. 7:**
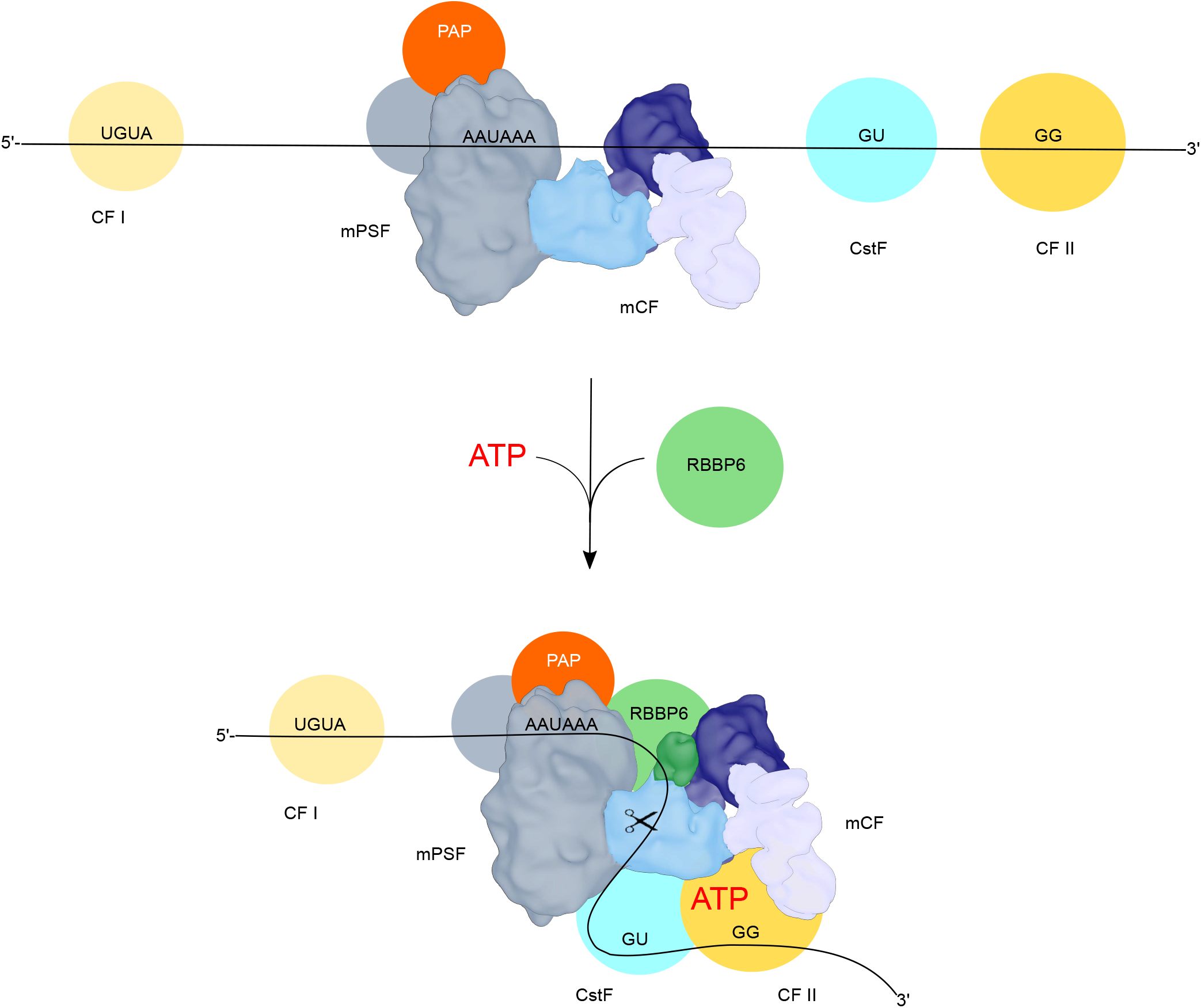
Model of the reconstituted pre-mRNA cleavage complex. RBBP6 and ATP are proposed to stabilize the activated conformation of the complex. CF I is not shown as part of the core complex, but it stimulates processing by interaction with hFip1 (Zhu et al. 2018).

In agreement with the original observations (Moore and Sharp 1985), pre-mRNA cleavage was found to depend on ATP but not cleavage of its phosphoanhydride bonds. ATP very likely serves as an allosteric effector for hClp1 since several mutations in the ATP binding site of this protein increase the concentration of ATP required for RNA cleavage, one mutation allows significant cleavage in the absence of ATP, and another mutation modulates the nucleotide specificity. An allosteric effect of ATP bound to hClp1 would be consistent with reports of mutations in the ATP binding site of yeast Clp1 affecting the interactions with other 3’ processing factors, notably Pcf11 (Holbein et al. 2011; Ghazy et al. 2012; Haddad et al. 2012). Still, all our hClp1 mutants could be purified in a stable complex with hPcf11. The yeast pre-mRNA 3’ cleavage reaction is independent of ATP addition (Hill et al. 2019). However, purified yClp1 contains tightly bound ATP (Noble et al. 2007; Holbein et al. 2011; Ghazy et al. 2012). Thus, a function of ATP in yeast pre-mRNA 3’ processing might be provided by the Clp1-bound nucleotide. As the K_A_ for ATP is ~0.5 μM, orders of magnitude below the typical intracellular concentration, the relevant ATP binding site in hClp1 will be occupied at all times, and variations in ATP binding are unlikely to serve a regulatory purpose. It is possible that ATP is a constitutive ingredient of the 3’ processing reaction with no regulatory significance. However, ATP hydrolysis, a reaction similar to the kinase activity of hClp1, might facilitate some aspect of the processing reaction that is not assayed in our experiments, in which non-hydrolyzable analogs were functional.

## Materials and Methods

### RNAs

The following RNAs have been described: SV40 late and SV40 lateΔ (Schäfer et al. 2018); L3 and L3Δ (Humphrey et al. 1987); ‘pre-cleaved’ RNA L3pre and L3preΔ (Christofori and Keller 1989). Mutations were introduced into L3 by means of synthetic DNA fragments corresponding to bp 4 - 369 in pSP64-L3 (Humphrey et al. 1987), counting from the transcription start site. One fragment contained G to C mutations in the UGUA-motifs at positions 144 and 155 (double mutant; GENWIZ), a second fragment contained an additional mutation at position 196 (triple mutant; Eurofins). Fragments were digested with BamHI/SacII and used to replace the corresponding fragment in pSP64-L3. The original L3pre RNA is lacking the UGUA upstream motifs. Therefore, L3pre-v2 was generated to include the UGUA motifs: Plasmids pSP64-L3 and pSP64-L3Δ as well as the derivatives carrying the UcUA mutations were used as templates for PCRs (for primers, see **Supplemental Table S6**) using Q5 DNA polymerase (NEB) to generate SP6 promotor-containing L3pre DNA fragments that end at the position of 3’-cleavage (after bp 198, counting from the transcription start, one nucleotide 3’ of the linearisation site in the original L3pre construct). The amplified DNA was purified, sequenced, and used for transcription. RNA was synthesized as described (Schäfer et al., 2018) in the presence of anti-reverse cap analog (NEB or Jena Biosciences) and gel-purified. A 5’- phosphorylated RNA of 51 nt corresponding to the expected SV40 late downstream cleavage fragment (Sheets et al. 1987) was synthesized by Biomers.net, Germany. The RNA was 3’ end-labeled with RNA ligase and [^32^P]-pCp.

### Processing assays

Reaction conditions for pre-mRNA cleavage underwent some evolution during this project. Under the most recent standard conditions, cleavage assays contained 10 μL 2x cleavage buffer (40 mM HEPES-KOH, pH 8.0, 4 mM DTT, 2 mM MgCl_2_, 1 mM ATP, 1 M trimethylamine oxide [TMAO]), 1 μg *E.coli* tRNA and up to 8 μL protein mix. The protein mix for each reaction contained 500 fmol each of CFI, CFII, CSTF, PAP D115A, mCF3; 60 fmol mPSF; 2500 fmol RBBP6_1-340_ or other RBBP6 variants as indicated; and 4 U murine RNase inhibitor (NEB). TMAO was adjusted to pH 7.5 in a 6 M stock solution with HCl. Proteins were pre-diluted to 1 μM in CDB200 (50 mM HEPES-KOH, pH 8.0, 20 % glycerol, 1 mM DTT, 0.5 mM EDTA, 200 mM KCL, 0.1 % Tween-20, 0.2 mg/ml RNase-free BSA [Merck]). The total reaction volume was made up to 20 μL with CDB200. In earlier assays, WT PAP was used in combination with 3’-dATP; 3’-dATP was also used in reactions with nuclear extract (non-dialyzed nuclear extract obtained from Ipracell, Mons, Belgium; 4 μl/reaction). Also, 4% polyvinyl alcohol (30 – 70 kDa) was initially used instead of TMAO and, with an older preparation, mPSF at 500 fmol/assay. Mixtures were assembled on ice and reactions started by the addition of 50 fmol substrate RNA and transfer to 30°C. After 1 h, they were stopped by the addition of SDS-containing buffer and proteinase K. After digestion, RNA was ethanol-precipitated and analyzed by 6% polyacrylamide-urea gel electrophoresis and phosphoimaging. For the calculation of cleavage efficiency, substrate RNA, 5’ cleavage fragment and background of each lane were quantified with ImageQuant 5.0. The background values were subtracted from the remaining substrate RNA and 5’ fragment, and the amount of 5’ fragment was corrected for the loss of radioactivity of the 3’ fragment. Percent cleavage was calculated by the division of the corrected 5’ fragment through the sum of 5’ fragment and uncleaved substrate RNA. For ATP titrations, data for a single time point were fitted to the Michaelis-Menten equation (SigmaPlot 12.5).

For RNase H/oligo(dT) digestion, purified RNA was dissolved in 3 μl RNase H reaction mix (1x RNase H buffer [NEB], 40 pmol oligo(dT)_12_, 4U murine RNase inhibitor [NEB], 1U RNase H [NEB]) and incubated for 30 min at 37°C. Reactions were stopped by addition of 3 μl formamide loading buffer and analyzed on denaturating polyacrylamide gels.

For the analysis of the 3’ cleavage fragment by primer extension, a 3x standard cleavage reaction was set up except that unlabeled SV40 late RNA was used and the amounts of this RNA and mPSF were doubled. RNAs were purified by phenol - chloroform extraction and ethanol precipitation, dissolved in 3 μl H_2_O, and 1 μl 0.5 μM primer (5’ ^32^P- labelled and gel-purified; CCCCCTGAACCTGAAACATA] and 1 μl dNTPs (10 mM each) were added. Mixtures were incubated for 5 min at 65°C and slowly cooled to room-temperature. 5 μl RT mastermix (2x reaction buffer, 20 mM DTT, 4U murine RNase inhibitor [NEB], 30 U ProtoScript II Reverse Transcriptase [NEB]) were added, and reactions were incubated for 1 h at 42°C and then stopped with 10 μl formamide loading buffer. RNAs were separated on a denaturing 10% polyacrylamide gel.

Polyadenylation reactions were done as described (Schönemann et al. 2014).

### RNA binding assays

Nitrocellulose filter binding was done in a 40 μl volume as described (Schäfer et al. 2018) except that the buffer was 20 mM HEPES-KOH, pH 7.9, 100 mM NaCl, 1 mM MgCl_2_, 0.5 mM ATP, 0.04 % Tween 20, 2 mM DTT. Proteins were pre-diluted in 50 mM HEPES-KOH, pH 7.9, 100 mM NaCl, 10 % glycerol, 0.5 mM EDTA, 0.05 % Tween 20, 0.2 mg/ml BSA, 1 mM DTT. Affinity measurements were carried out with 0.1 nM SV40 late RNA. Higher RNA concentrations were used to follow RNA binding activity across column fractions. For K_D_ determination, the binding equilibrium was expressed as a quadratic equation, and data were fitted in Microsoft Excel Solver.

### RNA kinase assays

Assays were performed according to (Schäfer et al. 2018) except that NP40 in the reaction buffer was replaced by Tween-20 and reactions contained 1 μM C_14_ RNA (3’-labeled by ligation to [5’-^32^P]-pCp and gel-purified), 1 nM CFII and 0.5 mM nucleotide. The reactions were stopped by the addition of 1 volume 8 M Urea and analyzed on a denaturing 9% polyacrylamide gel.

### Protein-protein interaction assays

For pull-down assays with purified proteins, anti-FLAG M2 antibody (Sigma F1365) was added to magnetic Protein G Dynabeads (Thermo Fisher) equilibrated in wash buffer (20 mM HEPES-KOH pH 7.9, 50 mM NaCl, 20 mM KCl, 5 mM MgCl_2_, 0.01% NP-40 substitute). After rotation at 4°C for 30 min, beads were washed three times and used immediately. 25 pmol FLAG-tagged RBBP6 was mixed with an equal amount of putative binding partner (final concentration 1 μM each) and incubated for 60 min at 4°C in 20 mM HEPES-KOH pH 7.9, 55 mM NaCl, 1 mM MgCl_2_, 20 mM creatine phosphate, 0.4 mM DTT. Equilibrated beads were added and rotated for 60 min at 4°C. The beads were washed three times with 20 volumes of wash buffer and bound proteins eluted with 0.2 mg/mL 3x FLAG peptide in wash buffer.

For co-precipitation assays using HEK cell lysates, 5×10^6^ cells per condition were transiently transfected using polyethyleneimine, induced immediately with doxycycline and harvested after 48 h. Flash-frozen cells were thawed and lysed in wash buffer as above (with 0.1% NP-40 substitute), and EDTA-free cOmplete Protease Inhibitor Cocktail (Roche) was added. Automated pull-down experiments were performed using magnetic Streptactin-coupled Dynabeads M270 (Thermo Fisher) and a KingFisher pull-down system (Thermo Fisher) operated at room temperature. After binding for 30 min, beads were washed four times and bound proteins eluted in SDS-containing sample buffer.

### Cryo-EM sample preparation, data collection and data processing

Before grid preparation, RBBP6 was run over a Superdex 200i 3.2/300 column (Cytiva) in 20 mM HEPES-KOH pH 7.9, 50 mM NaCl, 20 mM KCl, 5 mM MgCl_2_. The peak fraction was mixed with an equal amount of CPSF (final concentration at 1.5 μM) in the same buffer and incubated for 40 min at 4°C. 0.04 % (v/v) n-octyl-ß-D-glucoside was added directly before 4 μL of sample were applied onto glow-discharged Quantifoil R1.2/1.3, Cu 200 mesh grids. Grids were blotted for 3.5 s and plunge-frozen in a liquid ethane/propane mix using a Vitrobot Mark IV (Thermo Fisher) operated at 4°C and 100 % humidity.

Cryo-EM data were collected on a FEI Titan Krios microscope (Thermo Fisher) equipped with a post-column GIF (energy width 10 eV) and a Gatan K3 camera used in counting mode. The nominal magnification during data collection was 81,000x, corresponding to a pixel size of 1.094 Å at the specimen level. Using a beam-tilt based acquisition scheme in SerialEM (Schorb et al. 2019), the sample was imaged with a total exposure of 63 e^−^/Å^2^ evenly spread over 9.5 s and 63 frames. The target defocus ranged between −0.7 and −2.8 μm.

Movies were pre-processed on-the-fly using Focus (Biyani et al. 2017) while automatically discarding images of poor quality. Picked candidate particles were extracted in Relion 3.1 (Zivanov et al. 2018). After several rounds of reference-free 2D classification, particles were imported into CryoSPARC v3.1 (Punjani et al. 2017) for further processing in 3D (**Suppl. Fig. 4; Suppl. Table 7**). Structure visualization, analysis and rigid-body model docking was carried out using UCSF ChimeraX v1.2.5 (Pettersen et al. 2021) and PyMOL v2.3.2.

Expression constructs, cell culture conditions, protein purification, and mass-spectrometry are described in the Supplement.

## Supporting information

Complete Supplement

## Acknowledgements

We thank Anne-Katrin Hoffmeister and Gudrun Scholz for technical assistance; Erik Fiedler for suggesting the use of TMAO; David Bentley, Walter Keller, David G. Skalnik and Steven West for cDNAs; Daniel Bollschweiler and Tillman Schäfer at the MPIB cryo-EM facility for help with EM data collection; Barbara Steigenberger at the MPIB biochemistry core facility for mass spectrometry; Christian Benda and J. Rajan Prabu for providing computational infrastructure for EM data processing; Daniela Wartini for help with cell culture; Fabien Bonneau, Courtney Long and members of the Conti lab for discussions. This work was supported by a grant from the German Research Foundation to E.W. and by funding from the Max Planck Gesellschaft, the European Commission (ERC Advanced Investigator Grant EXORICO), the Novo Nordisk Foundation (Exo-Adapt) and the German Research Foundation (DFG Sonderforschungsbereich 1035, Projektnummer 201302640 and SFB/TRR 237) to E.C.

## Author contributions

F.S., U.K. and E.W. conceived the study. M.S., F.K., F.S., P.S., C.T., C.I., and U.K. performed experiments. M.S., F.K., F.S., U.K., E.C. and E. W. wrote the manuscript. All authors discussed and commented on the manuscript.

