## Supplementary material for "Reconstitution of 3’-processing of mammalian pre-mRNA reveals a central role of RBBP6": Complete Supplement

#### Supplemental Methods

##### Expression clones

Cloning mostly followed standard ligation-dependent procedures using enzymes from NEB. Reverse transcription of human ORFs was done using purified total RNA from HEK293 cells as described (Schönemann et al. 2014). New flanking restrictions sites were introduced by PCR using proof-reading polymerases (Pwo, Peqlab; Q5, NEB) and appropriate DNA primers (Invitrogen). Point mutations were introduced by the QuickChange protocol (Agilent) or as described by (Picard et al. 1994). All PCR products were verified by DNA sequencing (Eurofins). In some cases, ORFs were subcloned into additional vectors for fusion with N-terminal peptide tags or additional restriction sites before transfer into the MultiBac system. Alternatively, peptide tags were introduced into plasmids by ligation of phosphorylated oligonucleotides with compatible overhangs.

Plasmids and protocols for the MultiBac system were those described in (Fitzgerald et al. 2006; Bieniossek et al. 2008). Plasmids and protocols for the BigBac system are described in (Weissmann et al. 2016; Weissmann and Peters 2018) (Addgene). Individual ORFs were integrated into plasmids either by restriction digestion and ligation or by Gibson assembly (NEB). If more than two expression cassettes were combined in plasmids, this was performed either using DNA ligase and restrictions sites in the multiplication modules of the MultiBac plasmids or plasmids were fused by loxP-dependent Cre recombinase (NEB) (Bieniossek et al. 2008). The transfer vector for the PP1alpha complex was generated by Gibson assembly of four gene cassettes and the SwaI-opened pBIG1 vector according to (Weissmann and Peters 2018) with minor variations.

All cDNAs and proteins used in this study are listed in **Supplemental Table 3**. Affinity tags and other additional amino acid sequences are reported in **Supplemental Table 4**. Oligonucleotides used for mutagenesis and reverse transcription are listed in **Supplemental**

**Table 6**. Detailed information about the strategy used for cloning of individual open reading frames as well as DNA oligonucleotides used will be provided on request.

**Cell culture and virus propagation**

Suspension cultures of Sf21 cells (Invitrogen) were grown at 27.5°C on a rotary shaker in ExCell 420 serum-free medium (Sigma Aldrich) and diluted to 0.5 x $10^6$ cells/ml every 2 – 3 days. Cells were transfected with bacmids either with Cellfectin II (Invitrogen), and the supernatant (virus P1) was harvested after 3 days, or with Fugene HD (Promega), and the supernatant was harvested after 5 days. For virus propagation, cells were infected with 0.02 to 0.1 % culture volume of P1 and cultivated for up to 7 days. For protein expression, cells were infected at 1.5 x $10^6$ cells/ml with 1 % culture volume of the propagated virus and harvested 68 to 72 h later. Viruses for protein production are listed in **Supplemental Table 5.**

**Protein purification**

All proteins used for the reconstitution reaction were human except where noted. Procedures for the expression and purification of the following proteins have been described: CF I (Schäfer et al. 2018), full-length bovine PAP and bovine PABPN1 (Kühn et al. 2009). Bovine $PAP_{1-513}$ was purified according to (Martin and Keller 1996) except that a Resource Q column was used for the second step. CF II was purified as described (Schäfer et al. 2018) except that the final gel filtration column was omitted. All protein purifications described below were carried out under refrigeration. The pH of buffers was adjusted at room temperature in 1 M stock solutions. For Ni-NTA chromatography, the pH was re-adjusted after refrigeration in the final buffer. Ni-NTA beads were from Qiagen, anti-FLAG M2 agarose from Sigma, and all other column materials and pre-packed columns from Cytiva. Sf21 cells were harvested by centrifugation, resuspended in the buffer specified for each protein, and lysed by sonification with a Branson Sonifier, typically with 5 x 20 bursts at medium setting. Bacteria were lysed with a Constant Systems (Daventry, UK) device. Lysates were centrifuged for 30 min at 20,000 g.

Purification of mPSF was modified from (Schönemann et al. 2014). Cells from a 4 l culture were resuspended in 120 ml mPSF buffer (50 mM TEA pH 8.0, 10% sucrose, 0,5 mM DTT, 0.2 mM AEBSF, 2 µg/ml leupeptin, 2 µg/ml pepstatin) containing 250 mM KCl and lysed as described above. The cleared lysate was adjusted to pH 8.0 by the addition of KOH and loaded on a 30 ml DEAE-Sepharose CL6-B column equilibrated in mPSF buffer plus 250 mM KCl. The flow-through was agitated overnight with 0.3 ml Ni-NTA-agarose equilibrated in mPSF buffer plus 250 mM KCl, 5 mM imidazole. Beads were washed with 3 x 3 ml mPSF buffer plus 250 mM KCl, 10 mM imidazole, and protein was eluted with 4 x 3 ml of the same buffer plus 500 mM imidazole. The first three fractions were pooled, dialysed for 2 x 1 h against mPSF buffer plus 250 mM KCl and bound to 1 ml anti-FLAG beads for 2 h with agitation. Beads were washed with 10 ml mPSF buffer plus 250 mM KCl, and protein was eluted with 3 x 0.5 – 1 ml of the same buffer containing 0.15 mg/ml FLAG peptide (Sigma). Fractions were pooled and dialysed for 2 x 1.5 h against mPSF buffer plus 250 mM KCl containing 25 mM HEPES-KOH, pH 8.0, instead of TEA.

CstF was purified from 800 ml overexpressing Sf21 cells by a modification of (Schäfer et al. 2018). Cells were resuspended in 50 ml CstF buffer (50 mM TEA pH 8.0, 10% sucrose, 1 mM DTT, 1 mM EDTA, 1 mM PMSF) plus 200 mM KCl, 10 mM imidazole, and lysed as described above. The cleared lysate was agitated with 3 ml Ni-NTA-agarose. Beads were washed for 15 minutes with 10 column volumes of CstF buffer plus 200 mM KCl, 25 mM imidazole. Protein was eluted for 2 x 20 - 30 minutes with 15 ml CstF buffer plus 200 mM KCl, 250 mM imidazole. Elution fractions were pooled, diluted to 75 mM KCl with CstF buffer, and loaded on a 1 ml Mono Q column. The column was eluted with gradients of 20 column volumes from 75 to 500 mM KCl and of 5 column volumes to 1 M KCl, both in CstF buffer.

CstFΔ50 complexes were expressed in 400 ml Sf21 cells. Cells were resuspended in CstFΔ buffer (50 mM Tris pH 8.0, 10 mM imidazole, 10% glycerol, 0.02% NP-40, 1 mM DTT, 1 mM PMSF, 1 µg/ml leupeptin and 1 µg/ml pepstatin) plus 300 mM KCl and 10 mM imidazole. Cleared lysates were agitated with 300 µl Ni-NTA beads. Beads were washed three times with 10 column volumes of CstFΔ buffer plus 300 mM KCl and 10 mM imidazole. Protein was eluted with five times 2 column volumes CstFΔ buffer plus 200 mM KCl and 500 mM imidazole. Elution fractions were pooled, dialysed against CstFΔ buffer plus 75 mM KCl, and loaded on a 1 ml Resource Q column. The column was washed with the same buffer. Proteins were eluted with gradients of 15 column volumes from 75 to 500 mM KCl and 5 column volumes to 1 M KCl, all in CstFΔ buffer.

Sf21 cells expressing mCF3 or mCF4 were resuspended in mCF buffer A (50 mM Tris-HCl pH 7.5, 10 % sucrose, 5 µM $ZnCl_2$, 5 mM DTT, 1 mM PMSF, 1 µg/ml leupeptin and 1 µg/ml pepstatin) plus 75 mM KCl and lysed as described above. The cleared lysate was loaded on a 30 ml DEAE-Sepharose column. The column was washed with 2 column volumes mCF buffer A plus 75 mM KCl, and proteins were eluted with a three column volume gradient to 600 mM KCl and a final step of 1 M KCl, both in mCF buffer A. Fractions containing mCF were pooled, dialysed against mCF buffer B (20 mM HEPES-KOH, pH 7.9, 10 % sucrose, 5 µM $ZnCl_2$ and 2.5 mM DTT) plus 75 mM KCl and loaded on a 6 ml Resource S column. The flow-through was loaded on two combined 5 ml HiTrap Heparin-Sepharose columns in mCF buffer B plus 75 mM KCl. Proteins were eluted with a fifteen column volume gradient to 400 mM KCl and a final step of 1 M KCl in mCF buffer B. Fractions containing mCF were pooled and precipitated for 30 min on ice with ammonium sulphate at 50% saturation. The precipitate was collected by centrifugation (20 min, 20,000 g), dissolved in mCF buffer B plus 200 mM KCl with a reduced DTT content (1 mM), and proteins were separated on a 49 ml Superose 6 column in the same buffer.

Sf21 cells expressing CDC73 were resuspended in CDC73 buffer (50 mM potassium phosphate, pH 7.5, 10 % glycerol, 1 mM DTT, 1 mM PMSF, 1 µg/ml leupeptin and 1 µg/ml pepstatin) plus 300 mM KCl, 10 mM imidazole, and lysed as described above. The cleared lysate was agitated for 2 h with 0.5 ml Ni-NTA agarose. Beads were washed twice with 10 column volumes CDC73 buffer plus 300 mM KCl, 10 mM imidazole, and the protein was eluted with CDC73 buffer plus 300 mM KCl, 500 mM imidazole. 950 µl of the main elution fraction was diluted with CDC73 buffer to 100 mM KCl and loaded on a 1 ml Resource Q column. The column was washed with 2 column volumes CDC73 buffer plus 100 mM KCl and eluted with a 15 column volume gradient to 1 M KCl in CDC73 buffer.

E. coli BL21-CodonPlus expressing his-tagged SUMO-Ssu72 were grown in SB-medium at 37°C. Expression was induced with 500 µM IPTG at $OD_{600}$ = 1 and incubation continued overnight at 16°C. Cells were harvested, resuspended in 500 mM NaCl, 100 mM Tris-HCl, pH 8.0, and frozen in liquid nitrogen. Purification on a Ni-NTA column was carried out as described (Zhang et al. 2011). Elution fractions containing the Ssu72 fusion protein were pooled, mixed with his-tagged SUMO-protease, and dialyzed against 100 mM NaCl, 20 mM Tris-HCl, pH 8.0, 10 mM β-mercaptoethanol. The dialysate was agitated for 2 h with Ni-NTA agarose. The flow-through containing SUMO-free Ssu72 was collected.

Sf21 cells expressing RBBP6 or its variants were resuspended in RBBP6 buffer (50 mM Tris/HCl pH 8.0, 10 % sucrose, 1 mM DTT, 10 µM $ZnCl_2$,1 mM PMSF, 1 µg/ml pepstatin, 1 µg/ml leupeptin) plus 200 mM KCl, 10 mM imidazole and lysed as described above. Cleared lysates were bound for 2 – 4 h to Ni-NTA-Agarose beads equilibrated in the same buffer. The beads were washed with the same buffer, and protein was eluted with the same buffer containing 250 mM imidazole. For further purification of the deletion variants, relevant elution fractions were pooled, dialysed against RBBP6 buffer containing 50 mM KCl, centrifuged and loaded on a Resource Q column. The column was eluted with three consecutive gradients: 2 column volumes to 200 mM KCl, 10 column volumes to 400 mM KCl and 3 column volumes to 1 M KCl, all in RBBP6 buffer. Full-length constructs tagged with MBP, MBP-SNAP, or GFP as well as the His-FLAG-tagged ΔRING mutant were pooled after Ni-NTA purification, diluted to a final concentration of 150 mM KCl with RBBP6 buffer, and loaded on a Resource Q column. The column was eluted with a ten column volume gradient up to 400 mM KCl and then three column volumes up to 1 M KCl. MBP and GFP tags were cleaved with PreScission protease and the MBP-SNAP tag with TEV-protease. Proteases were used at a 1:50 ratio for two hours at room temperature. *E. coli* BL21-CodonPlus cells expressing shortened RBBP6 variants were harvested, resuspended in RBBP6 buffer plus 200 mM KCl and lysed. Purification was as described for proteins expressed in Sf21 cells except that 10 % glycerol replaced sucrose.

Sf21 cells expressing XRN2 were resuspended in XRN2 buffer (50 mM Tris/HCl pH 7.5, 10 % sucrose, 3 mM $MgCl_2$, 1 mM DTT, 10 µM $ZnCl_2$, 1 mM PMSF, 1 µg/ml pepstatin 1 µg/ml leupeptin) containing 200 mM KCl, 10 mM imidazole, and lysed as described above. The protein was purified on a Ni-NTA column as described for RBBP6. Elution fractions containing XRN2 were pooled, dialysed against XRN2 buffer plus 75 mM KCl, and loaded on a Resource Q column. The protein was eluted with two gradients of 15 column volumes to 500 mM KCl and 3 column volumes to 1 M KCl in the same buffer.

Purification of the PP1α complex from Sf21 cells was carried out as described for full-length Rbbp6 except that Tris-HCl was used at pH 7.5 and wash buffer for the Ni-NTA column contained 20 mM imidazole.

The RNA polymerase II CTD with an N-terminal GST fusion was expressed in *E. coli* BL21-CodonPlus or Sf21 cells. Bacteria were harvested, resuspended in 100 mM KCl, 50 mM Tris-HCl pH 7.5, 10% glycerol, 1 mM DTT, 0.5 mM PMSF, 0.8 µg/ml pepstatin, 0.5 µg/ml leupeptin and lysed. Sf21 cells were resuspended in the same buffer and lysed. In both cases, cleared lysates were bound to equilibrated GSH-Sepharose. Elution was carried out in lysis buffer plus 10 mM reduced glutathione. Fractions containing the desired protein were loaded on a Superdex 200 HR10/30 size exclusion column. The column was run in lysis buffer containing 5 mM DTT and lacking glutathione. For dephosphorylation, 2 µg of a fraction from the final Superdex column was incubated with 2.5 units Quick CIP (NEB) at 37°C. Aliquots were withdrawn at different time points and analysed by SDS polyacrylamide gel electrophoresis and Coomassie staining or by western blotting with an antibody specific for Ser2P (CTD7 3E10) (Chapman et al. 2007).

The concentration of $RBBP6_{1-340}$ was determined in a spectrophotometer at 280 nm. Concentrations of all other proteins were determined by densitometry (ImageQuant 5.0; Molecular Dynamics) of Coomassie R250-stained bands in SDS polyacrylamide gels and comparison to BSA standards.

**Mammalian cell protein expression and purification**

HEK293T stable expression cell lines were established using the *piggybac* transposon system by initially transfecting the cells using polyethyleneimine (Yusa et al. 2011; Li et al. 2013). Pools of cells were generated that stably expressed either RBBP6 (RBBP6 residues 1-335 N-terminally fused to a TwinStrep tag and a 3C protease cleavage site), mCF4 (TwinStrep-3C-Symplekin, CPSF100, CPSF73, CstF64), or CPSF (TwinStrep-3C-Symplekin, CPSF100, CPSF73, CstF64, CPSF160, WDR33 residues 1-413, CPSF30). Additionally, RBBP6 variants carrying an N-terminal TwinStrep-3C tag and a C-terminal FLAG tag (RBBP6 residues 1-335, wildtype and mutant forms) were transiently expressed using polyethyleneimine transfection.

For protein expression, cultures were adjusted to a density of $1x10^6$ cells per mL in FreeStyle 293 expression medium (Gibco, Thermo Fisher). The cells were induced with doxycycline and harvested 48 h after induction. For all purifications, lysis buffers were supplemented with EDTA-free complete Protease Inhibitor Cocktail (Roche), DNase I and Benzonase before cells were lysed with a glass dounce homogenizer and cleared by centrifugation.

All RBBP6 variants were purified in 1x DPBS supplemented with 2 mM DTT, 10 µM $ZnCl_2$. Cleared lysate was applied onto a StrepTrap HP column, washed with 20 column volumes of buffer and eluted using 5 mM desthiobiotin in wash buffer. The N-terminal TwinStrep tag was removed by addition of 3C protease and the target protein was further purified over a MonoQ 5/50 GL column. After washing, bound protein was eluted using a gradient increasing the salt concentration to 1000 mM NaCl. Fractions containing pure protein were dialyzed overnight into 1x DPBS supplemented with 2 mM DTT, 10 µM $ZnCl_2$, 2 mM $MgCl_2$, concentrated and flash frozen in liquid nitrogen.

The four-subunit mCF was purified in 1x DPBS supplemented with 2 mM DTT, 10 µM $ZnCl_2$ using a StrepTrap HP column. After washing, bound proteins were eluted with 5 mM desthiobiotin in wash buffer, concentrated and further purified over a Superose 6i 10/300 gel filtration column. Fractions containing pure protein were pooled, concentrated and flash frozen in liquid nitrogen.

Seven-subunit CPSF was purified in 1x DPBS supplemented with 2 mM DTT, 10 µM $ZnCl_2$, 2 mM $MgCl_2$ using a StrepTrap HP column. After extensive washing, bound proteins were eluted with 5 mM desthiobiotin in wash buffer. Protein preparations used for cryo-EM were concentrated and flash frozen in liquid nitrogen. All other protein preparations were applied to a HiTrap Heparin column for further purification. After washing, bound proteins were eluted using a gradient increasing the salt concentration to 1000 mM NaCl. Fractions containing pure protein were dialyzed overnight into 1x DPBS supplemented with 2 mM DTT, 10 µM $ZnCl_2$, 2 mM $MgCl_2$, concentrated and flash frozen in liquid nitrogen.

**Cross-linking and mass spectrometry**

For cross-linking mass spectrometry with BS3, 1.0 µM CPSF and RBBP6 were mixed with an RNA substrate in a buffer containing 20 mM HEPES-KOH pH 7.9, 55 mM NaCl and 1 mM $MgCl_2$. The sample was incubated 45 min at 4˚C before 0.5 mM BS3 was added. After 20 min incubation at 4˚C, the reaction was quenched by adding ~ 40 mM Tris-HCl pH 7.5 and incubating 15 min at 4˚C. The sample was spun for 10 min at 18,000 g. For denaturation of the crosslinked proteins, 4 M Urea and 50 mM Tris was added to the supernatant and the samples were sonicated using a Bioruptor Plus sonication system (Diogenode) for 10x 30 sec at high intensity. For reduction and alkylation of the proteins, 40 mM 2-cloroacetamide (CAA, Sigma-Aldrich) and 10 mM tris(2-carboxyethyl)phosphine (TCEP; Thermo Fisher) were added. After incubation for 20 min at 37°C, the samples were diluted 1:2 with MS grade water (VWR). Proteins were digested overnight at 37°C by addition of 1 µg of trypsin (Promega). Thereafter, the solution was acidified with trifluoroacetic acid (TFA; Merck) to a final concentration of 1%, followed by desalting of the peptides using Sep-Pak C18 1cc vacuum cartridges (Waters). The elution was vacuum dried and the desalted peptides were further pre-fractionated into 8 fractions using a high pH reversed-phased nano-fractionation system (Kulak et al. 2017).

Fractionated peptides were loaded onto a 30-cm analytical column (inner diameter: 75 microns; packed in-house with ReproSil-Pur C18-AQ 1.9-micron beads, Dr. Maisch GmbH) by the Thermo Easy-nLC 1000 (Thermo Fisher) with buffer A (0.1% (v/v) formic acid) at 400 nl/min. The analytical column was heated to 60°C. Using the nanoelectrospray interface, eluting peptides were sprayed into the benchtop Orbitrap Q Exactive HF (Thermo Fisher Scientific) (Hosp et al. 2015). As gradient, the following steps were programmed with increasing addition of buffer B (80% acetonitrile, 0.1% formic acid): linear increase from 8 to 30% over 60 minutes, followed by a linear increase to 60% over 5 minutes, a linear increase to 95% over the next 5 minutes, and finally maintenance at 95% for another 5 minutes. The mass spectrometer was operated in data-dependent mode with survey scans from m/z 300 to 1650 Th (resolution of 60k at m/z = 200 Th), and up to 15 of the most abundant precursors were selected and fragmented using stepped Higher-energy C-trap Dissociation (HCD with a normalized collision energy of value of 19, 27, 35). The MS2 spectra were recorded with dynamic m/z range (resolution of 30k at m/z = 200 Th). AGC target for MS1 and MS2 scans were set to 3 x $10^6$ and $10^5$, respectively, within a maximum injection time of 100 and 60 ms for the MS1 and MS2 scans, respectively. Charge state 2 was excluded from fragmentation to enrich the fragmentation scans for cross-linked peptide precursors. The acquired raw data were processed using Proteome Discoverer (version 2.5.0.400) with the XlinkX/PD nodes integrated (Klykov et al. 2018). To identify the crosslinked peptide pairs, a database search was performed against a FASTA containing the sequences of the proteins under investigation. DSS was set as a crosslinker. Cysteine carbamidomethylation was set as fixed modification and methionine oxidation and protein N-term acetylation were set as dynamic modifications. Trypsin/P was specified as protease and up to two missed cleavages were allowed. Furthermore, identifications were only accepted with a minimal score of 40 and a minimal delta score of 4. Otherwise, standard settings were applied. Filtering at 1% false discovery rate (FDR) at peptide level was applied through the XlinkX Validator node with setting simple.

For crosslinking of the pre-mRNA cleavage complex with DSBU, PAP and RBBP6 were dialyzed against 25 mM HEPES-KOH, pH 8.0, 200 mM KCl, 10 % sucrose, 1 mM DTT to remove Tris buffer; all other proteins were present in buffers lacking amines. Reactions with or without PVA were set up. Each reaction contained 250 µL 2x cleavage buffer (see above), 6.25 pmol SV40 late RNA, 25 µg *E.coli* tRNA, 50 µl CDB200, 12.5 pmol each of CFI, CFII,

CstF, bPAP, mCF3, mPSF and either 62.5 pmol RBBP6$_{1-340}$ or 31.25 pmol RBBP6$_{1-780}$. The reaction volume was increased to 500µl with 25 mM HEPES-KOH, pH 8.0, 200 mM KCl, 10 % sucrose, 1 mM DTT. After pre-incubation for 30 minutes at 30°C, 1.5 µl DSBU (Thermo Fisher; 50 mM in DMSO) was added and the incubation continued for another 30 minutes at 25°C. The reaction was stopped by the addition of 25 µl 1 M Tris-HCl, pH 8.0. Proteins were precipitated by the addition of TCA to 10% and pelleted by centrifugation. Pellets were washed with 80 % ice-cold acetone, air-dried, and dissolved in 25 µl 8 M urea in 0.4 M ammonium bicarbonate. Sulfide bonds were reduced for 20 min at 50°C with 7.5 mM DTT, and alkylation was performed at 25°C for 20 min with 14 mM chloroacetamide. The volume was made up to 200 µl with $H_2O$, and 2.5 µg trypsin (sequencing grade modified trypsin, Promega) was added. After incubation over night at 37°C, the reaction was stopped by the addition of 20 µl 10 % TFA. Peptides were first purified by StageTips (Rappsilber et al. 2007). Per reaction 10 StageTips with 5 Empore $C_{18}$ extraction discs (Sigma-Aldrich) were prepared and washed with 20 µl methanol, then with buffer B (60% acetonitrile, 0.1% TFA) and finally buffer A (0.1% TFA). Equal volumes of the crosslink reactions were loaded on the StageTips. The StageTips were washed with 20 µl buffer A and eluted with 5 µl buffer B. The eluate was loaded on a Superdex Peptide 3.2/300 column (GE Healthcare) running at 25 µL/min in 3% acetonitrile, 0.1% TFA. Fractions of 50µl were collected and analyzed by nano-HPLC/nano-ESI-MS/MS using an Ultimate 3000 RSLC nano-HPLC system coupled to an Orbitrap Q-Exactive Plus mass spectrometer equipped with NanoFlex source (both Thermo Fisher). Samples were injected on a trap column (PepMap RP-C18, 300 µm × 5 mm, 5 µm, 100 Å, Thermo Fisher) with 0.1 % TFA at a flow rate of 30 µL/min. After 15 min, peptides were eluted via a 200 cm µPAC™ separation column (Pharmafluidics). For separation, 240 min gradients were applied (3-6.5 % B over 25 min at 600 nL/min followed by 6.5-33 % B over 205 min at 300 nL/min; A: 0.1 % formic acid in water, B: 0.08 % formic acid in acetonitrile). Data were acquired in data-dependent MS/MS mode (one MS survey scan, followed by 10 MS/MS scans of the 10 most abundant signals). MS survey scans (*m/z* range 375-1799) were recorded with $R$ = 140000 at *m/z* 200, the target of the automated gain control (AGC) was set to $3x10^6$ and the maximum injection time was 100 ms. For MS/MS scans, triply to seven times protonated ions were selected for fragmentation by HCD with stepped normalized collision energies (NCEs: 27, 30, and 33 V). The following parameters were applied for MS/MS scans: $R$ = 17.500, AGC: 2 × $10^5$ and maximum injection time: 250 ms; dynamic exclusion was set to 60 s. The raw files were converted to mgf files by MsConvert (Chambers et al. 2012). Crosslinks were analyzed by Merox v. 2.0.1.4 (Gotze et al. 2015) and visualized with the help of xView (Graham et al. 2019).

**Supplemental Table 1: Comparison of 3' end processing factors in mammals and in *S. cerevisiae***

| name | subunits | *S. cerevisiae* orthologue | Yeast orthologue part of |
|---|---|---|---|
| mPSF | CPSF160 (CPSF1)<br>WDR33<br>hFip1 (FIP1L1)<br>CPSF30 (CPSF4) | Cft1<br>Pfs2<br>Fip1<br>Yth1 | CPF (polymerase module)<br>CPF (polymerase module)<br>CPF (polymerase module)<br>CPF (polymerase module) |
| mCF | CPSF100 (CPSF2)<br>CPSF73 (CPSF3)<br>symplekin | Cft2<br>Ysh1<br>Pta1 | CPF (nuclease module)<br>CPF (nuclease module)<br>CPF (phosphatase module) |
| CF I | CF I-59 (CPSF7)<br>or CF I-68 (CPSF6)<br>CF I-25 (CPSF5, NUDT21) | -<br>-<br>- | |
| CF II | hPcf11<br>hClp1 | Pcf11<br>Clp1 | CF IA<br>CF IA |
| CstF | CstF77 (CSTF3)<br>CstF64 (CSTF2)<br>or CstF64$\tau$ (CSTF2T)<br>CstF50 (CSTF1) | Rna14<br>Rna15<br>Rna15<br>- | CF IA<br>CF IA<br>CFIA |
| RBBP6 | | Mpe1 | CPF (nuclease module) |
| PAP | | PAP | CPF (polymerase module) |
| PABPN1 | | - | |
| - | | Hrp1 | CF IB |

For a description of yeast CPF and its three modules and of CF IA and CF IB see (Casanal et al. 2017).

**Supplemental Table 2: Apparent affinities of 3' processing factors for RNA**

| Factor | $K_{50}$ (nM) |
|---|---|
| mPSF | 0.7; 0.8 |
| mCF3 | 1.6; 3.3; 3.8 |
| mCF4 | 0.8; 0.9; 2.1; 4.4 |
| CF I | 210; 212 |
| CF II | 4; 6 |
| $RBBP6_{1-780}$ | 55; 67 |
| CstF (64/64) | 7; 8 |
| CstF ($\tau/\tau$) | 18; 23 |
| CstF (64/$\tau$) | 15 |

Apparent affinities were measured by nitrocellulose filter-binding experiments using the SV40 late RNA (see Methods). Data listed represent the results of individual titration experiments.

**Supplemental Table 3: Proteins/ORFs used in this study.**

| ORF/protein | UniProt/Genbank | citation /source |
|---|---|---|
| hFip1 | Q6UN15 | (Kaufmann et al. 2004; Schönemann et al. 2014) |
| hWDR33 (WDC146) | Q9C0J8 | (Ito et al. 2001; Schönemann et al. 2014) |
| hSymplekin | Q92797 | (Schönemann et al. 2014) |
| hCPSF1 (=CPSF160) | Q10570 | (Schönemann et al. 2014) |
| hCPSF4 (=CPSF30 | O95639-2 | (Schönemann et al. 2014) |
| hCPSF2 (=CPSF100) | Q9P2I0 | SinoBiological |
| hCPSF3 (=CPSF73) | Q9UKF6 | SinoBiological |
| hRbbp6 | Q7Z6E9 | Promega (#FHC25637) |
| eYFP | - | (Berger et al. 2004) |
| CFP (cerulean protein) | CAO79586.1 (gb) | (Schönemann et al. 2014) |
| mCherry | ADO95149.1 (gk) | (Schönemann et al. 2014) |
| hPcf11 | O94913 | (Schäfer et al. 2018) |
| hClp1 | Q92989 | (de Vries et al. 2000; Schäfer et al. 2018) |
| CSTF1 (= CstF50) | Q05048 | (Schäfer et al. 2018) |
| CSTF2 (= CstF64) | P33240 | (Schäfer et al. 2018) |
| CSTF2$\tau$ (=CstF64$\tau$) | Q9H0L4 | Walter Keller |
| CSTF3 (= CstF77) | Q12996 | (Dettwiler et al. 2004; Schäfer et al. 2018) |
| CPSF5 (= CF I-25) | O43809 | (Dettwiler et al. 2004; Schäfer et al. 2018) |
| CPSF6 (= CF I-68) | Q16630 | (Dettwiler et al. 2004; Schäfer et al. 2018) |
| bovine PAP alpha | P25500 | (Kühn et al. 2017) |
| bovine PABPN1 | Q28165 (NP-776994) | (Kühn et al. 2017) |
| mouse RNA Pol II CTD | P08775 | David Bentley |
| PP1 complex: | | |

| | | |
|---|---|---|
| hPP1$\alpha$ | P62136 | Sino Biological (HG10533-M), David G. Skalnik |
| mouse Pnuts | Q80W00 (mouse) | David G. Skalnik |
| hWDR82 | Q6UXN9 | this work |
| hTox4 | O94842 | David G. Skalnik |
| hSsu72 | Q9NP77 | Promega (# FHC08360) |
| hCDC73 | Q6P1J9 | this work |
| hXRN2 and XRN2 D235A | Q9H0D6 | Steven West |

### order number

#### Supplemental Table 4: Affinity-tags and additional amino acids in recombinant proteins of this study

| Protein | N-terminal peptide fusions and additional amino acids |
|---|---|
| CFP | MVLA – CFP |
| Cherry | MVLA – mCherry |
| eYFP | eYFP – RSI |
| His-CF I-25 (CPSF5, NUDT21) | MSHHHHHHHHGDP – CF I-25 (aa2-227) |
| Strep-CF I-68 (CPSF6) | MWSHPQFEKG – CF I-68 |
| His-Clp1<br>and mutants E22A, E22Q or R288A/R293L | MAHHHHHH – Clp1 |
| Strep-Pcf11<br>and variants | MWSHPQFEKARGA – Pcf11 |
| Flag-CPSF160 (CPSF1) | MDYKDDDDKAIA – CPSF160 |
| CPSF160 (CPSF1) | CPSF160 – V |
| CPSF100 (CPSF2) | none |
| CPSF73 (CPSF3) | none |
| Strep-CPSF73 (CPSF3) | MWSHPQFEKLA – CPSF73 |
| CPSF30 (CPSF4) | none |
| Fip1 | none |
| Myc-His-WDR33 | MEQKLISEEDLHHHHHHTS – WDR33 |
| Symplekin | none |
| TwinStrep-3C-Symplekin | MSAWSHPQFEKGGGSGGGSGGSAWSHPQFEKTAGLEVLFQGP - Symplekin |
| Strep-CstF50 (CSTF1) | MWSHPQFEKDP – CstF50 (aa2-431) |
| CstF64 (CSTF2) | None |
| Flag-CstF64τ (CSTF2τ) | MDYKDDDDKLD – CstF64tau |
| His-CstF77 (CSTF3) | MSHHHHHHHHGDP – CstF77 (aa2-717) |
| His-Rbbp6-Flag | MSHHHHHHHHAIA – Rbbp6 – VLDYKDDDDK |
| HF-Rbbp6<br>and C-terminal deletion variants | MSHHHHHHHHADYKDDDDKAIA – Rbbp6 |
| HF-Rbbp6-ΔRING | MSHHHHHHHHADYKDDDDKAIA – hRbbp6 (aa1-252 + aa327-1792) |
| His-MBP-Rbbp6 | MGSSHHHHHHSSGR – MBP – NSSSNNNNNNNNNNSSGRLEVLFQGPAIA – Rbbp6<br>GPAIA – Rbbp6 (after HRV 3 C cleavage) |
| RBBP6-FLAG aa1-335 (HEK cells) and variants | GP - RBBP6 (aa1-335) - DYKDDDDK |

| | |
|---|---|
| pBIG1a-PP1A complex:<br>His-WDR82 | <br>MSHHHHHHHHGDP – WDR82 |
| Tox4 | none |
| PP1a | none |
| mouse Flag-mPnuts | MDYKDDDDKASVVEF – mPnuts |
| His-CDC73 | MSHHHHHHHHHQF – CDC73 |
| His-SUMO-Ssu72 | RDL – Ssu72 (after cleavage) |
| His-Xrn2 and mutant D235A | MSHHHHHHHHASRASEA – Xrn2 |
| GST-mouse RNA Pol II CTD | MTGGQQ – GST – SDLVPRGSV – mCTD (aa1586-1970/S1724 is missing) |
| bovine PAP alpha and variants | MAHHHHHH – bPAP |
| bovine His-PABPN1 | MAHHHHHH – bPABPN1 |

N- or C-terminal extensions are listed in single letter code. MBP - maltose binding protein; CFP – Cerulean/cyan fluorescent protein

**Supplemental table 5: Viruses used for recombinant expressions**

| Virus /Bacmid | vectors used for Bacmid | fused via |
|---|---|---|
| pBac-mPSF$^{Tn7}$ | pKL- FlagCPSF1 – CPSF4<br>pUCDM-HisMyc-WDR33 – hFip1 healed | Cre/loxP<br>recombinase |
| pBac untagged mCF3$^{loxP}$<br><br>and substitution CPSF3 mutant:<br>D75K/H76A | pSPL-CFP – Symplekin – CPSF3-CPSF2<br><br>in the same construct context | multiplication<br>module |
| pBac untagged mCF3$^{loxP}$ – CstF2 $^{Tn7}$<br>(mCF4) | pSPL-CFP – Symplekin – CPSF3 – CPSF2<br>pFBDM-CstF2 | |
| pBAC-mCherry$^{loxP}$ – CstF $^{Tn7}$ | pUCDM-mCherry<br>pFBDM-StrepCstF1 – CstF2 – HisCstF3 | multiplication module |
| pBAC-mCherry$^{loxP}$ – CstF$\tau$ $^{Tn7}$ | pUCDM-mCherry<br>pKL-HisCstF3 – FlagCstF2 tau – StrepCstF1 | multiplication module |
| pBAC-mCherry$^{loxP}$ – (CstF+ $\tau$)$^{Tn7}$ | pUCDM-mCherry<br>pKL-His-CstF3-Flag-CstF2 tau-StrepCstF1 – CstF2 | multiplication module |
| pBAC-mCherry$^{loxP}$ – CstF$\Delta$50 $^{Tn7}$ | pUCDM-mCherry<br>pFBDM-CstF2 – HisCstF3 | |
| pBAC-mCherry$^{loxP}$ – CstF$\tau$ $\Delta$50 $^{Tn7}$ | pUCDM-mCherry<br>pFBDM-FlagCstF2 tau – HisCstF3 | |
| pBAC-mCherry$^{loxP}$ –CstF1 $^{Tn7}$ | pUCDM-mCherry<br>pFBDM-Strep-CstF1 | |
| pBAC-(YFP) $^{loxP}$ -<br>(CPSF5-CPSF6) $^{Tn7}$ | pUCDM-YFP polH<br>pKL-HisCPSF5 – StrepCPSF6 | |
| pBAC-(YFP) $^{loxP}$ -(hPcf11-hClp1) $^{Tn7}$<br>and hClp1 substitution variants: E22A, E22Q, R288A/R293L | pUCDM-YFP<br>pFL– HisClp1 – StrepPcf11 or<br>in the same context as above | |
| pBAC-mcherry$^{loxP}$ – His-Rbbp6-CFlag $^{Tn7}$ | pUCDM-mCherry<br>pFBDM-His-Rbbp6-CFlag | |

| | | |
|---|---|---|
| pBAC-mcherry$^{loxP}$ – MBP-Rbbp6 $^{Tn7}$ | pUCDM-mCherry<br>pLIB-His-MBP-Rbbp6 | |
| pBac–mcherry$^{loxP}$–HF-Rbbp6 ΔU-box$^{Tn7}$ | pUCDM-mCherry<br>pLIB-HisFlag-Rbbp6 ΔU-box | |
| pBAC-mcherry$^{loxP}$ – (HF-Rbbp6 C-terminal deletions: 1-253, 1-340, 1-490, 1-780) $^{Tn7}$<br>and point mutations in 1-780:<br>C161G/C164G and N312D | pUCDM-mCherry<br>pLIB-HisFlag-Rbbp6 truncations<br><br>same context | |
| pBAC-mcherry$^{loxP}$ – GST-mCTD) $^{Tn7}$ | pUCDM-mCherry<br>pLIB-His-GST-CTD | |
| pBAC-YFP$^{loxP}$ His-Xrn2 wt$^{Tn7}$<br>and variant D235A | pUCDM-YFP<br>pFBDM-His-Xrn2 | |
| pBAC-YFP$^{loxP}$ – Cdc73$^{Tn7}$ | pUCDM-YFP<br>pFL-His-Cdc73 | |
| pBAC-mcherry$^{loxP}$ – PP1 complex $^{Tn7}$ | pUCDM-mCherry<br>pBIG1a-– His-WDR82– hTox4– hPP1α– Flag-mPnuts | Gibson assembly |

**Supplemental Table 6: Oligonucleotides used for mutagenesis and amplification of human cDNAs**

| usage | primer | sequence |
|---|---|---|
| CPSF3 D75K/H76A | 198 for | CATTTTATTTACAATCACTCGACG |
| | 342 rev | GTACACTGCCATACACTTCTTG |
| | 344 rev | AGAAACCAGGGCAAAGCTCCACAAGCTTTCAAATGGAAATGACTAATTAGTAGG |
| Clp1 mutagenesis | 684 int for | CCACCATCGGGCGCGGATCCAAGACTTGATCACCCGGGATCTC |
| | 685 int rev | TCCTCTAGTACTTCTCGACAAGCTTCCCCATCTCCCGGTACC |
| Clp1 E22A | 754a for | GAACTAGAGCGAGAAACAGCGCTTCGCTTTGAGGTGGAGGCA |
| Clp1 E22Q | 757 for | GAACTAGAGCGAGAAACACAACTTCGCTTTGAGGTGGAGGCA |
| Clp1 R288A/R293L | for | GGGTGTGGTGGAGGCCTCCAAGGACTTCCTACGTGAATGTAGGG |
| | rev | CCCTACATTCACGTAGGAAGTCCTTGGAGGCCTCCACCACACCC |
| Rbbp6 N312D | 723 for | CTCCTGATGCTTTAATTGCCGATAAGTTTTTACGACAGGCTGTAAATAAC |
| | 724 rev | GTTATTTACAGCCTGTCGTAAAAACTTATCGGCAATTAAAGCATCAGGAG |
| Rbbp6 C161G/C164G | 725 for | CCACCTCCATCTTACACGGGTTTCCGTGGTGGTAAACCTGGACATTATATTAAG |
| | 726 rev | CTTAATATAATGTCCAGGTTTACCACCACGGAAACCCGTGTAAGATGGAGGTGG |
| WDR82 cDNA | 366 5’UTR | TGGCCGACTTCCCGCTAG |
| | 367 3’UTR | CAGAAATAGCCAAGCAGCAAC |
| CDC73 cDNA | 382 for | GCTAGCGAATTCATGGCGGACGTGCTTAGCGT |
| | 383 rev | GGTACCAAGCTTTCAGAATCTCAAGTGCGATTTATGCTTTACC |
| L3pre UGUA double mutagenesis | 193 for | TCATATTGTCGTTAGAACGCG |
| | 744 for | TACAAATAAAAACATTTGCCT |
| L3pre UGUA triple mutagenesis | 745 rev | TAGAAATA-AAAACATTTGCCT |

#### Supplemental Table 7: Cryo-EM data collection and processing statistics

| **Data collection and processing** | |
|---|---|
| Microscope | FEI Titan Krios GII |
| Voltage (kV) | 300 |
| Camera | Gatan K3-Summit |
| Energy filter | Gatan Quantum-LS (GIF) |
| Magnification | 81,000x |
| Pixel size (Å/pix) | 1.094 |
| Target defocus range (µm) | -0.7 to -2.8 |
| Electron exposure ($e^-$/Å$^2$) | 63 |
| Number of movies | 7,462 |
| Initially selected particle candidates | 1,303,481 |
| Final number of particles | 69,204 |
| Resolution$_{\text{FSC independent halfmaps}}$ (Å)[a] | 7.98 |

[a] according to the Fourier Shell Correlation (FSC) cut-off criterion of 0.143 defined in (Rosenthal and Henderson 2003)

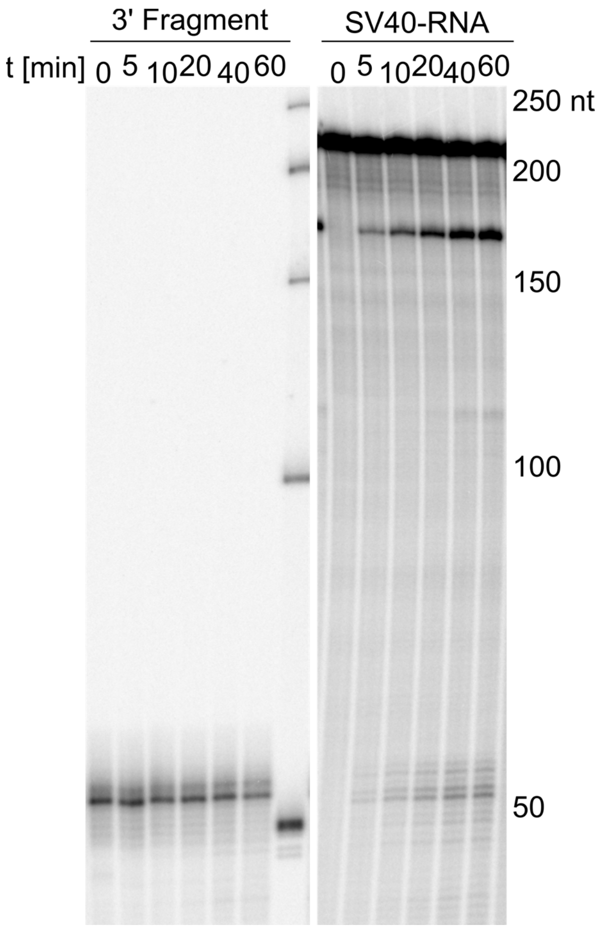


**Fig. S1: A synthetic downstream fragment is not degraded in the pre-mRNA cleavage reaction.**

Left panel: The 3’ end-labeled synthetic 3’ cleavage fragment (10 fmol) (see Methods) was incubated under standard cleavage conditions in the presence of all 3’ processing factors. Since the assay was part of an effort to investigate a possible function of XRN2, 500 fmol of the inactive XRN2 D235A mutant was also included. No degradation of the synthetic RNA was observed over the time course shown. Right panel: The SV40 late RNA was incubated under the same conditions to show that the reaction mixture was active in cleavage.

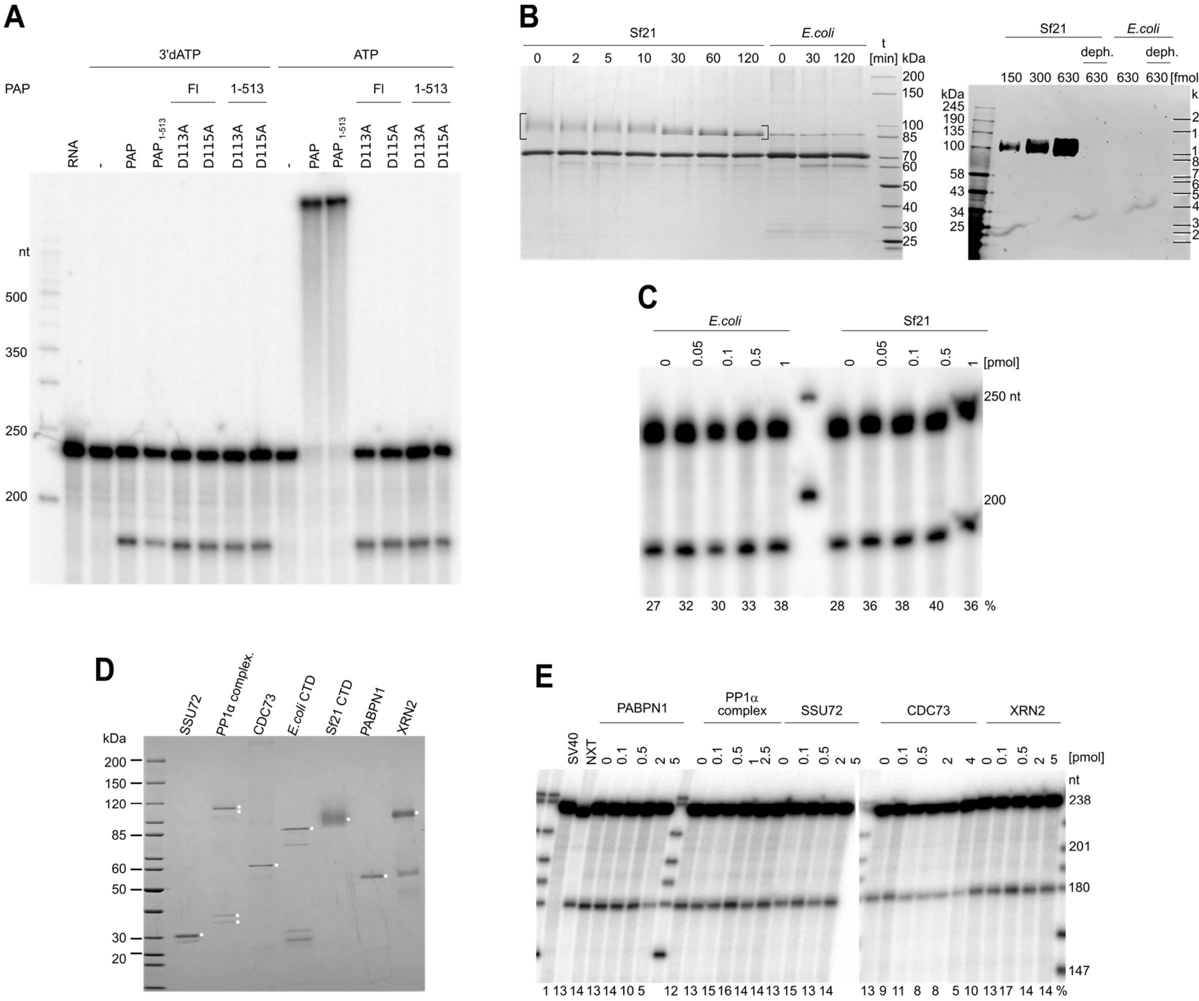


**Fig. S2: Roles of proteins in pre-mRNA cleavage.**

(A) Catalytic activity of poly(A) polymerase is not essential for RNA cleavage. Reconstituted cleavage reactions were carried out in the presence of PVA with different variants of PAP (full-length or C-terminally shortened $PAP_{1\text{-}513}$, either WT or with the point mutations indicated) and with either 3'dATP or ATP.

(B) The Sf21-derived CTD is Ser2-phosphorylated. Left panel: CTD prepared from Sf21 cells or *E. coli* was treated with phosphatase as described in Materials and Methods for the times indicated. Brackets indicate the heterogeneous migration of Sf21-derived CTD. Phosphatase treatment converted the Sf21-made protein to a homogeneous form co-migrating with the *E. coli*-made material, which was not affected by phosphatase treatment. Thus, the Sf21-derived protein is phosphorylated. The band below the CTD is BSA. Proteins were analyzed by SDS polyacrylamide gel electrophoresis and Coomassie staining. Right panel: CTD from Sf21 cells was recognized by an antibody specific for Ser2-phosphorylated CTD in western blots. Phosphatase-treated protein and E. coli-made CTD served as negative controls.

(C) The RNA polymerase II CTD has little effect on pre-mRNA cleavage. The CTD (mouse), made either in Sf21 cells or in *E. coli* as indicated, was titrated in pre-mRNA cleavage assays. Cleavage efficiencies are reported at the bottom. One representative experiment of two is shown.

(D) Coomassie-stained SDS gel showing additional purified proteins potentially involved in pre-mRNA cleavage. PABPN1 plays a role not only in polyadenylation (see Introduction) but also in alternative polyadenylation and thus cleavage site choice *in vivo* (de Klerk et al., 2012; Jenal et al., 2012). SSU72 is a known ligand of symplekin (Xiang et al., 2010) and the ortholog of the yeast phosphatase Ssu72, a subunit of the phosphatase module of CPF (Casanal et al., 2017). The protein phosphatase 1 (PP1) complex (Lee et al., 2010) is composed of PP1 (ortholog of yeast Glc7), PNUTS, TOX4, and WDR82, (ortholog of yeast Swd2). Like Ssu72, Glc7 and Swd2 are parts of the phosphatase module of CPF (Casanal et al., 2017). PP1 is thought to be involved in RNA polymerase II transcription termination (Cortazar et al., 2019; Kecman et al., 2018). CDC73 (parafibromin), a subunit of the Paf1 complex, has been reported to associate with CPSF and CstF and recruit the 3' processing complex to transcribed genes (Rozenblatt-Rosen et al., 2009). Finally, XRN2 is responsible for the degradation of the 3' cleavage product (Eaton et al., 2018; Kim et al., 2004; West et al., 2004). Desired polypeptides are labeled. In the PP1 complex, TOX4 and FLAG-PNUTS run as one band.

(E) Additional proteins do not stimulate pre-mRNA cleavage. Protein preparations shown in (D) were titrated in cleavage assays. Reactions contained 4% PVA and 500 fmol mPSF.

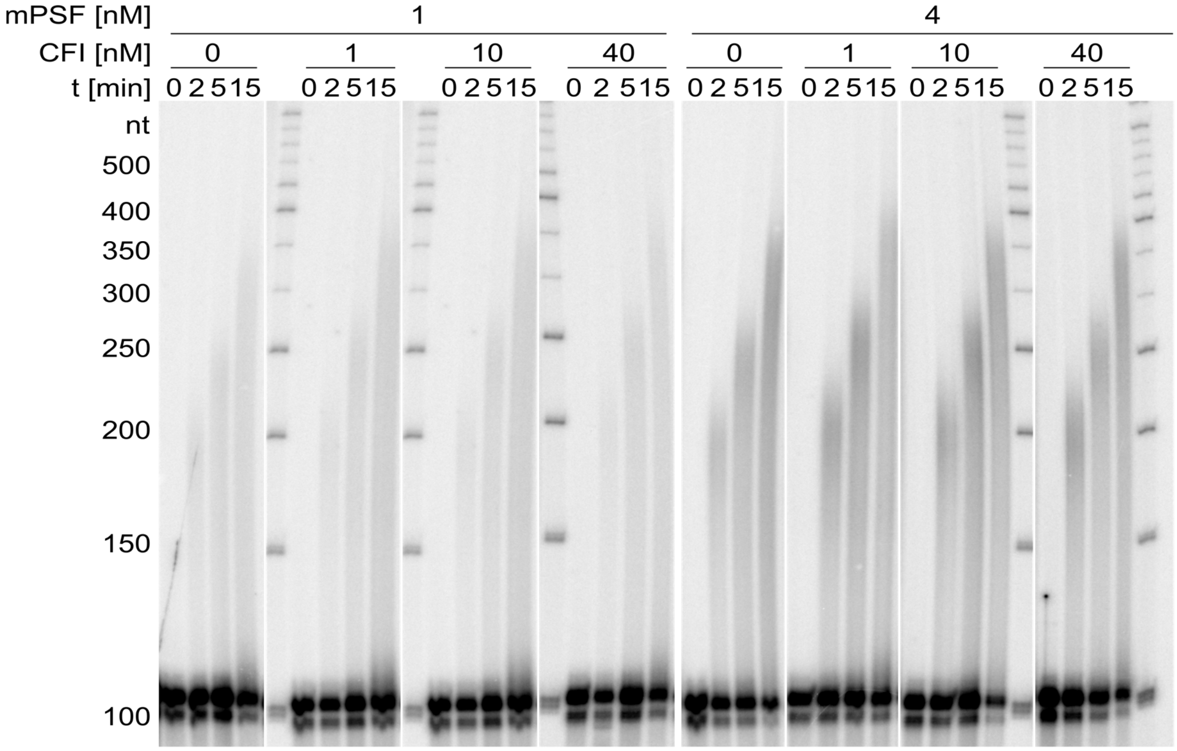


**Fig. S3: CF I does not stimulate a reconstituted polyadenylation reaction**

Polyadenylation time courses were carried out with 0.4 nM PAP and the concentrations of mPSF and CF I indicated. The substrate RNA (4 nM) was L3pre-v2 (see Materials and Methods).

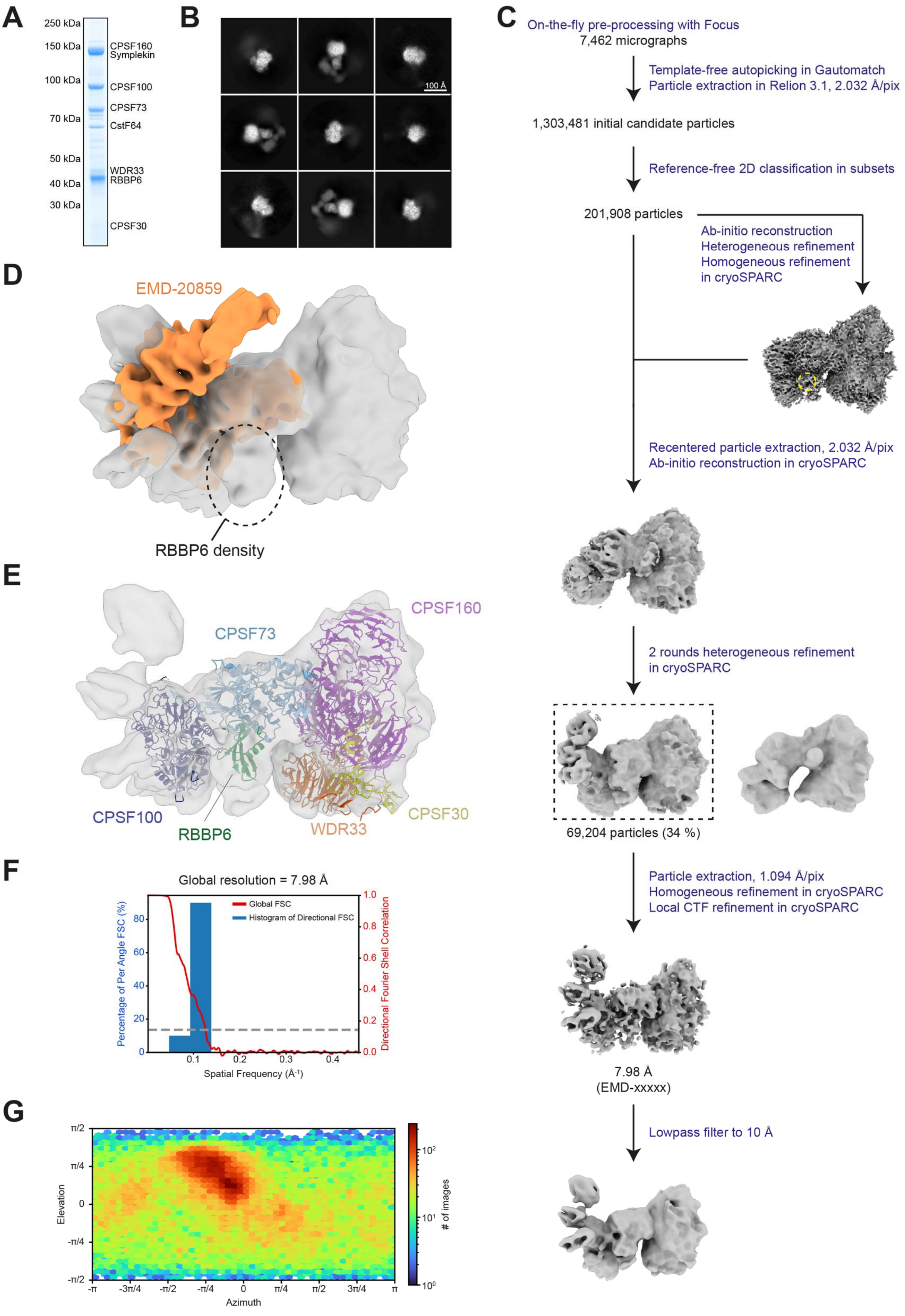

A
250 kDa
150 kDa
100 kDa
70 kDa
50 kDa
40 kDa
30 kDa
CPSF160
Symplekin
CPSF100
CPSF73
CstF64
WDR33
RBBP6
CPSF30
B
100 Å
C
On-the-fly pre-processing with Focus
7,462 micrographs
Template-free autopicking in Gautomatch
Particle extraction in Relion 3.1, 2.032 Å/pix
1,303,481 initial candidate particles
Reference-free 2D classification in subsets
201,908 particles
Ab-initio reconstruction
Heterogeneous refinement
Homogeneous refinement
in cryoSPARC
Recentered particle extraction, 2.032 Å/pix
Ab-initio reconstruction in cryoSPARC
2 rounds heterogeneous refinement
in cryoSPARC
69,204 particles (34 %)
Particle extraction, 1.094 Å/pix
Homogeneous refinement in cryoSPARC
Local CTF refinement in cryoSPARC
7.98 Å
(EMD-xxxxx)
Lowpass filter to 10 Å
D
EMD-20859
RBBP6 density
E
CPSF73
CPSF160
CPSF100
RBBP6
WDR33
CPSF30
F
Global resolution = 7.98 Å
Global FSC
Histogram of Directional FSC
Percentage of Per Angle FSC (%)
Directional Fourier Shell Correlation
Spatial Frequency (Å-1)
G
Elevation
Azimuth
### of images

**Fig. S4: Cryo-EM analysis of the CPSF-RBBP6 complex**

(A) Coomassie-stained SDS-PAGE showing the purified CPSF-RBBP6 complex.

(B) Representative 2D class averages of picked particles. Scale bar ~ 100 Å.

(C) Cryo-EM data processing scheme. Processing steps are indicated in blue; particle numbers of classes used for downstream processing and resolutions are in black. Yellow dashed circle indicates new center for particle extraction.

(D) Cryo-EM map of mCF alone (EMD-20859) in orange fitted into the cryo-EM map of the CPSF-RBBP6 complex. Both cryo-EM maps were filtered to 10 Å before superposition. The same orientation as in Fig. 4 (B) is shown and the extra density protruding from the proximal lobe is indicated by a black circle.

(E) Filtered cryo-EM map with fitted structural models of mPSF (PDB: 6fbs), CPSF100 (PDB: 6v4x), and CPSF73-RBBP6 (AlphaFold2 model). The same orientation as in Fig. 4 (B) is shown.

(F) Three-dimensional FSC plot (Tan et al., 2017): The red line represents the estimated global masked half map FSC. The resolution according to the gold standard FSC cut off of 0.143 is indicated and shown as grey dashed line (Rosenthal and Henderson, 2003).

(G) Plot visualizing the distribution of particle views.

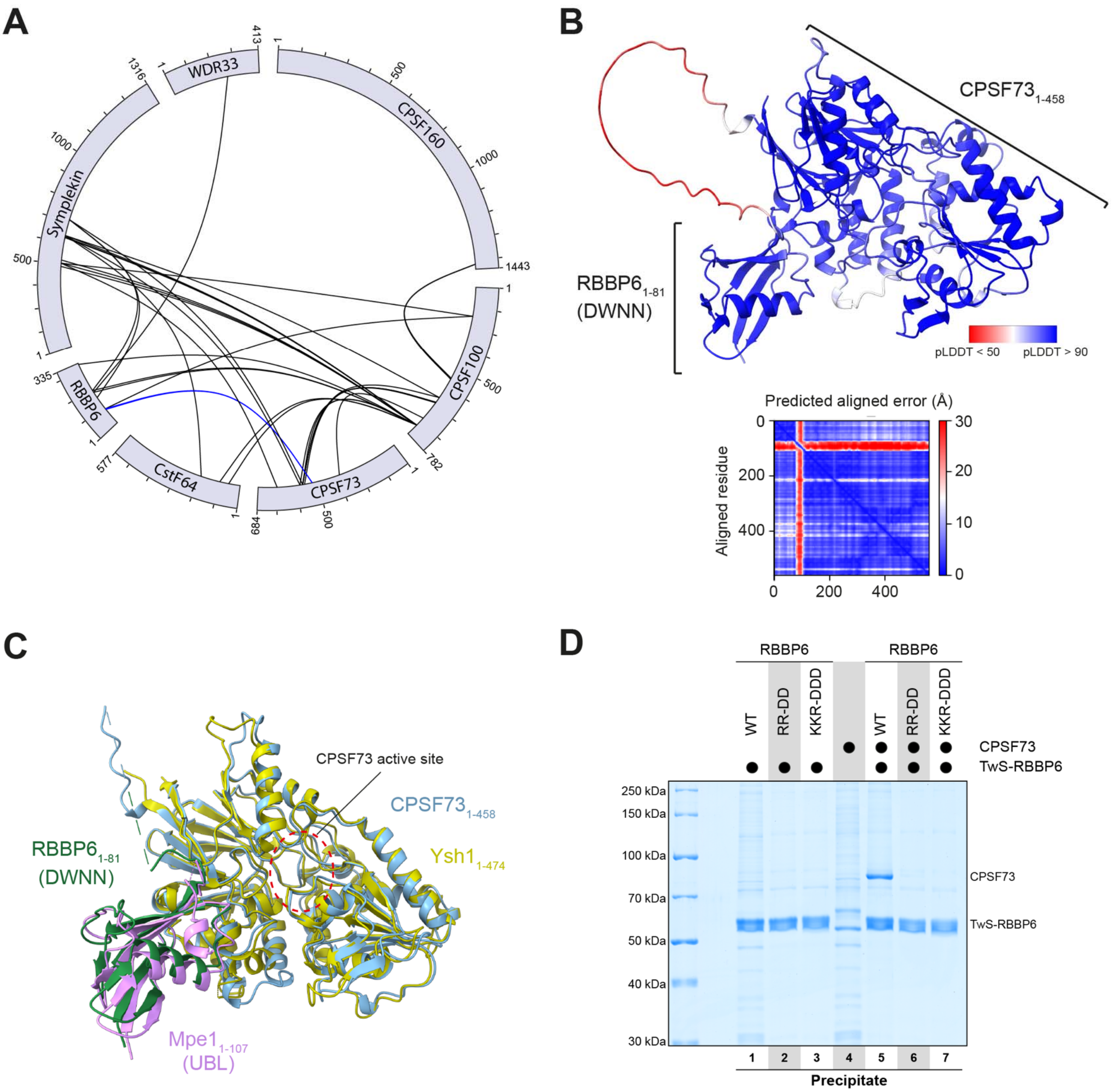


**Fig. S5: Characterization of the CPSF-RBBP6 interaction**

(A) BS3-dependent intermolecular cross-links of the CPSF-RBBP6 complex. Intermolecular cross-links between CPSF3 and RBBP6 shown in Fig. 4 (D) are highlighted in blue. Intramolecular cross-links are omitted for clarity.

(B) Computationally predicted structural model of the $CPSF73_{1-458}$-$RBBP6_{1-81}$ complex colored by its per-residue confidence score (pLDDT) (Jumper et al., 2021). The artificial linker sequence connecting the two domains is shown in red with low confidence score. Below, predicted aligned error plot with blue indicating high confidence in predicting relative positions.

(C) Overlay of the AlphaFold2 model of CPSF73-RBBP6 with experimentally derived structure of yeast Ysh1-Mpe1 (PDB: 6i1D) with an RMSD of 0.884 Å. Approximate location of the CPSF73 active site is indicated by a red dashed circle.

(D) Coomassie-stained SDS-PAGE analysis of pull-down experiment showing that the reverse-charged mutations in RBBP6 disrupt the interaction to CPSF73.

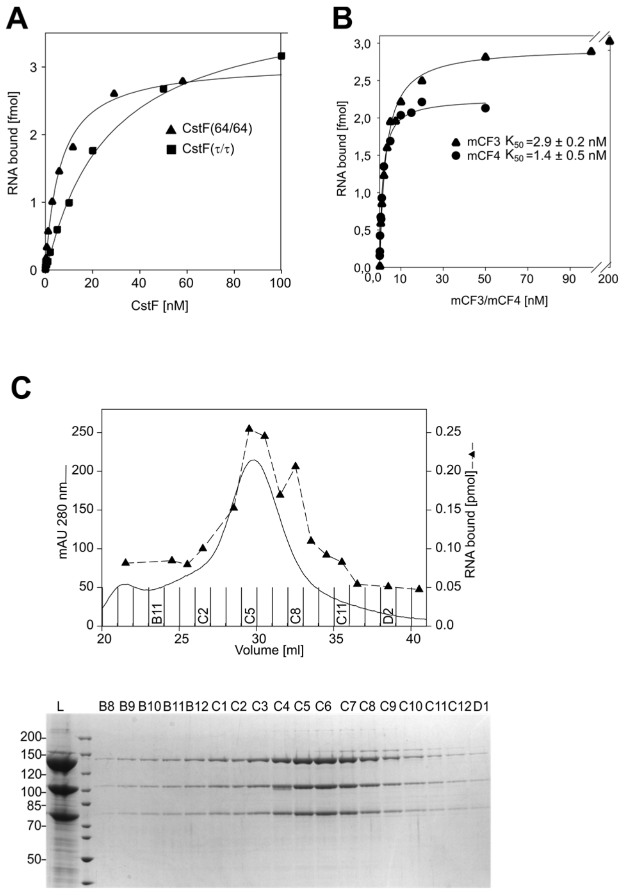


**Fig. S6: RNA binding activities of CstF and mCF**

(A) CstF (64/64) binds RNA with slightly higher affinity than CstF (τ/τ). Both proteins were titrated in nitrocellulose filter-binding assays. For each protein, data were averaged from two titrations. (B) mCF3 binds RNA with a similar affinity as mCF4. The proteins were titrated in filter-binding experiments with SV40 late RNA. The data are averaged from n = 3 (mCF3) or n = 4 (mCF4). (C) RNA-binding activity co-purifies with mCF3. The profile of the final Superose 6 column in the purification of mCF is shown. Upper panel: RNA binding activity in the column fractions was measured by nitrocellulose filter-binding assays and plotted over the UV profile. Lower panel: Aliquots from the column fractions were analyzed by SDS PAGE and Coomassie staining. L, load.

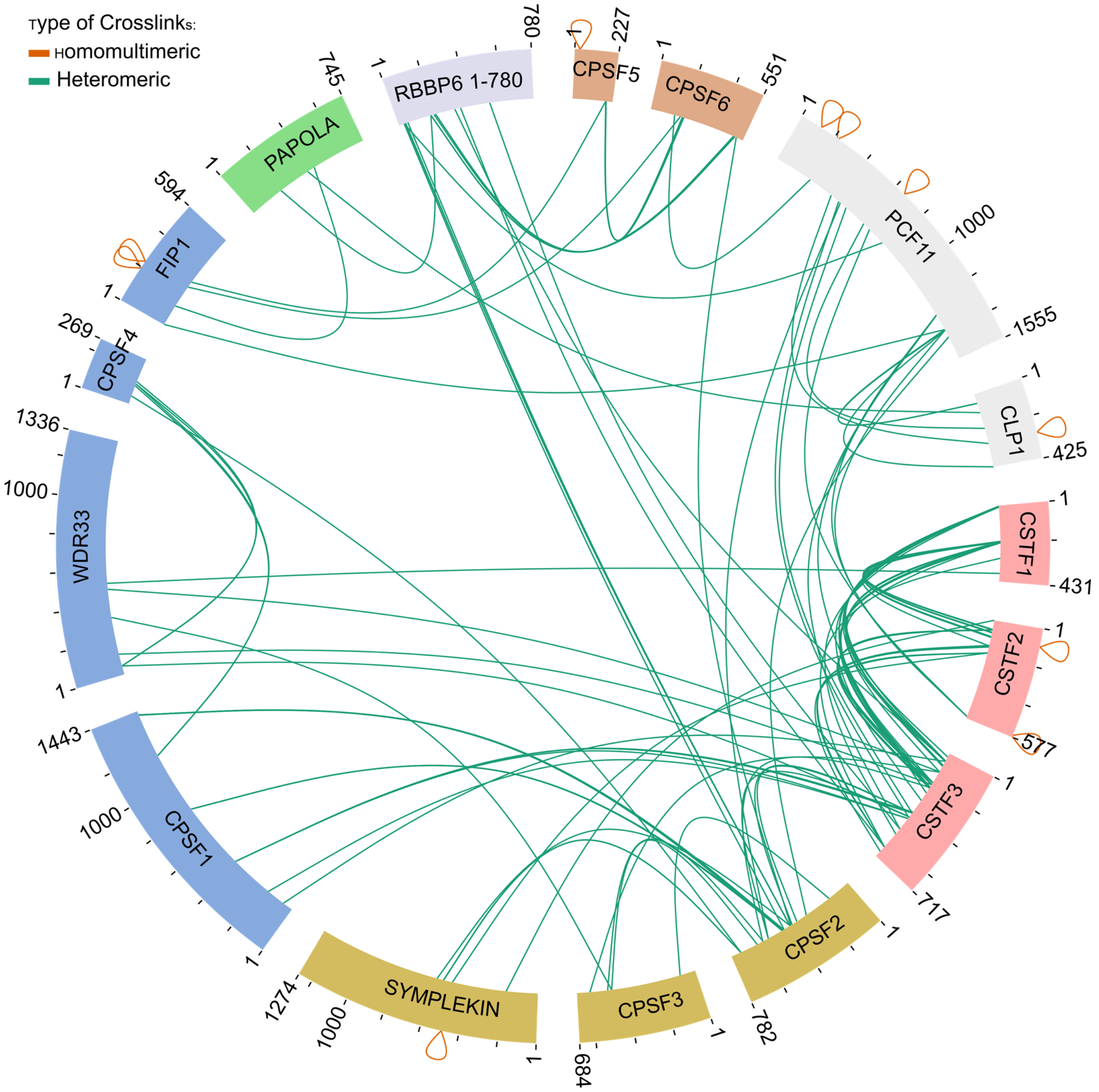


**Fig. S7: Summary of all DSBU-dependent intermolecular protein-protein cross-links**

Protein-protein cross-linking was carried out with DSBU in complete 3' processing reactions as described in Methods. The figure summarizes data from three independent experiments with and three without PVA. Red, cross-links that indicate homooligomerization of proteins.
